# Surface-stabilized sub-micron condensates for compartmentalizing synthetic cells and enhanced enzyme kinetics

**DOI:** 10.64898/2026.07.24.740618

**Authors:** Udit Ghosh, Elza van der Velde, Zohaib Hussain, Diane W. te Brake, Chang Chen, Chuanbao Zheng, Jasper van der Gucht, Renko de Vries, Siddharth Deshpande

## Abstract

Living cells harbor numerous membraneless organelles (MOs), which are dynamic protein/nucleic acid assemblies responding to specific physicochemical triggers. Liquid-liquid phase separation (LLPS) plays a crucial role in their formation and activity. Inspired by the natural MOs that maintain their individual identities, this work presents a bioengineering strategy to generate LLPS-driven, isochemical MO populations using surfactant-like peptides that stabilize the MO interface. The result is highly stable, monodispersed, sub-µm-sized MO populations, which are not only capable of compartmentalizing synthetic cells but also provide superior environments for enzymatic reactions. This is achieved using pH-responsive elastin-like polypeptides (PREs) as MOs and formulating an amphiphilic PRE-based peptide to stabilize the MO interface. Relative abundance of the surface-active peptide, as well as the rate of pH change, allows direct control over the MO size. Encapsulating these components within synthetic vesicles using a microfluidic platform leads to on-demand multi-compartmentalization via an external pH trigger. Lastly, a functional consequence of the acute size control is shown through a phosphatase reaction, where the highest reaction rate is observed in size-controlled MOs when compared to dilute environments and surfactant-free MOs. The presented strategy provides a new avenue for designing programmable MOs and thus achieve functional compartmentalization within synthetic cells.

## INTRODUCTION

Cells control biochemical activities by segregating them into different compartments^1^. While membrane-bound organelles, such as the nucleus, endoplasmic reticulum, Golgi bodies, and mitochondria, have long been known^2^, the past two decades have made it increasingly clear that cellular organization goes beyond mere membrane-bound organelles^3,4^. Membraneless organelles (MOs), also called biomolecular condensates, are usually mediated through liquid-liquid phase separation (LLPS) and play an important role in intracellular organization^5–7^. MOs can form because of weak and multivalent interactions, such as electrostatic, hydrophobic, hydrogen bonding, π-π, and cation-π interactions^8–11^. In contrast to membrane-bound organelles, MOs are highly dynamic and can assemble, reorganize, and disassemble depending on changes in the cellular environment^12–14^. Nucleoli provide an example of the ability of RNA-and protein-rich, liquid-like condensates to fuse and reorganize to generate larger compartments, where coalescence plays a role in determining the size and shape of nucleoli in *Xenopus laevis* oocytes^15^. NPM1-rich and FIB1-rich phases represent immiscible liquid-like subcompartments within nucleoli, where homotypic fusion and surface tension-driven organization lead to the generation of a multilayered nucleolar architecture^16^. In comparison, P granules in *C. elegans* are an example of a different organizational strategy. Instead of fusing to create a single body, P granules remain as several separate RNA-protein condensates, which are organized through regulated dissolution and condensation along the anterior-posterior axis, leading to the formation of posterior germline-associated P granules^17^. Overall, these examples indicate that LLPS can serve as a mechanism for organizing biological processes. However, since these are membraneless structures, they also pose challenges in preserving their distinct identities and preventing unwanted fusion with other MOs, whether of the same or different nature.

If their interface is not stabilized, liquid-like condensates tend to coalesce to lower their interfacial energy and undergo Ostwald ripening due to the variation in Laplace pressure between droplets of different sizes. However, cellular and engineered condensate systems often remain distinct through the regulation of their interfaces and material properties. Interfacial protein assemblies, ranging from endogenous MEG-3 clusters on P granules to engineered protein cages on reconstituted protein condensates, adsorb at the condensate interface and suppress coalescence and coarsening, thereby stabilizing the condensate population^18,19^. A related example outside MOs is Ki-67, which acts as a biological surfactant at the mitotic chromosome surface, keeping chromosomes dispersed rather than allowing them to collapse into a single chromatin mass^20^. In conventional polymer and colloidal systems, amphiphiles and diblock copolymers have been shown to self-assemble into mesoscale structures through the segregation of chemically dissimilar regions^21–23^. Additionally, minimal models show that surfactant-like client proteins are able to lower the surface tension of MOs enough to allow multiple condensates of stable size to form^24^. Recent studies have shown that amphiphilic block copolymers self-assemble at the interface between dense and dilute phases, generating an interfacial layer that preserves droplet fluidity and influences fusion, permeability, and interfacial mechanics^25–27^. Similar observations are now emerging in biological and biomimetic systems. Transcriptional regulators such as MLX, MYC, and IPMK act as surfactant-like proteins for TFEB MOs and lower interfacial tension, thereby reducing TFEB-DNA affinity and transcriptional activity^28^. Engineered amphiphilic proteins adsorb to the MO surface and control condensate size and shape, providing additional evidence that MO size can be controlled directly through the design of amphiphilic protein architecture^29,30^. Together, these examples show that condensate interfaces play an active role in controlling condensate growth, stability, and interactions with the surrounding phase.

Although substantial progress has been made in understanding how biomolecular condensates (MOs) stabilize their interfaces and regulate biochemical reactions, several unanswered questions remain. First, how can the size of an MO be precisely controlled across the mesoscale to engineer multicompartmentalized synthetic cells? Achieving such control is important for generating stable and reproducible internal compartments with defined sizes and numbers. Second, how do MOs regulate enzymatic reactions when the enzymes and substrates are not covalently coupled to the phase-separating scaffold but instead partition into the dense phase? Previous studies have often demonstrated kinetic enhancement within MOs using systems where there is a direct interaction between enzymes and MO scaffolds^31–34^. For example, MOs formed through the fusion of two distinct low-complexity domains to NADH oxidase showed catalytic enhancement owing to the local enrichment of substrates and cofactors, together with the physicochemical environment within the dense phase^35^. In another study involving lipases, dense phase formation created a less polar and locally buffered microenvironment that allowed reactions to be compartmentalized under different pH conditions, enabling cascade reactions between enzymes with contrasting pH optima^36^. More broadly, recent work on coacervate protocells has shown that phase-separated condensates can influence reaction kinetics through molecular partitioning, exchange between the dense and dilute phases, and changes in the local physicochemical environment^37^. However, whether such catalytic enhancement can also be achieved without covalent coupling between the enzyme and the MO scaffold remains unclear.

Here, we describe a simple yet effective strategy to control MO size and thus their number and assess the functionality of the resulting compartments through enzymatic reaction kinetics. Our system is based on elastin-like polypeptides (ELPs), a class of recombinant protein polymers inspired by elastin and characterized by repetitive pentapeptide motifs, (GXGVP)_n_, where X is a guest residue and n is the number of pentapeptide repeats^38–40^. The phase behaviour of ELPs depends on both sequence composition and polypeptide chain length^41,42^, and they typically exhibit lower critical solution temperature (LCST)-type LLPS that can be tuned by physicochemical parameters such as temperature, pH, salt conditions, and osmolyte concentration^43–46^. We use PREs that we previously engineered, allowing LLPS to be triggered by reducing the solution pH below the transition pH^47^. To address the size-control challenge, we developed an ELP-based amphiphilic diblock surfactant-like protein, Surf-PRE, that exhibits high affinity for the condensate interface, leading to the formation of monodisperse MO populations, from µm-sized down to 200 nm. Confocal imaging demonstrated the localization of the surfactant at the surface of MOs, while dynamic light scattering (DLS) analysis indicated the long-term stability of the MO populations. We further employ on-chip microfluidics to generate water-in-oil-in-water double emulsion vesicles as our cell-mimetic system^48,49^. By encapsulating the PRE along with its surfactant counterpart, we demonstrate stable formation of multiple MOs within each synthetic cell, where the size and the number of MOs are dictated by the surfactant-like protein as well as the acidification rate. Finally, using a phosphatase reaction, we show that these MOs enhance enzymatic reaction rates even without direct coupling between the enzyme and the phase-separating scaffold, displaying superior reaction kinetics, with the highest reaction rate observed in size-arrested MOs.

## RESULTS

### Designing ELP-based amphiphilic surfactants to stabilize the MO interface

As the MO-forming material, we used PRE-*h*-36, hereafter simply referred to as PRE (Figure 1a). PRE consists of the ELP sequence (GXGVP)_60_, where X represents valine, tyrosine, glutamic acid, or histidine in a 7:8:4:1 ratio (Table S1). PRE undergoes LLPS below its transition pH of 5.7 at room temperature, leading to the formation of PRE-rich MOs. The PRE-encoding plasmid was verified by Sanger sequencing using the primers listed in Table S2, after which PRE was expressed and purified by inverse transition cycling (ITC; see Methods). The molecular weight of PRE and Surf-PRE was further confirmed by SDS-PAGE and matrix-assisted laser desorption/ionization time-of-flight mass spectrometry (MALDI-TOF; Supplementary Figure S1. To visualize PRE-rich MOs by fluorescence microscopy, the terminal cysteine residue of PRE was conjugated to Alexa Fluor 647 (AF647)-maleimide. Unless otherwise stated, AF647-PRE constituted 4 mol% of the final PRE concentration.

**Figure 1.**
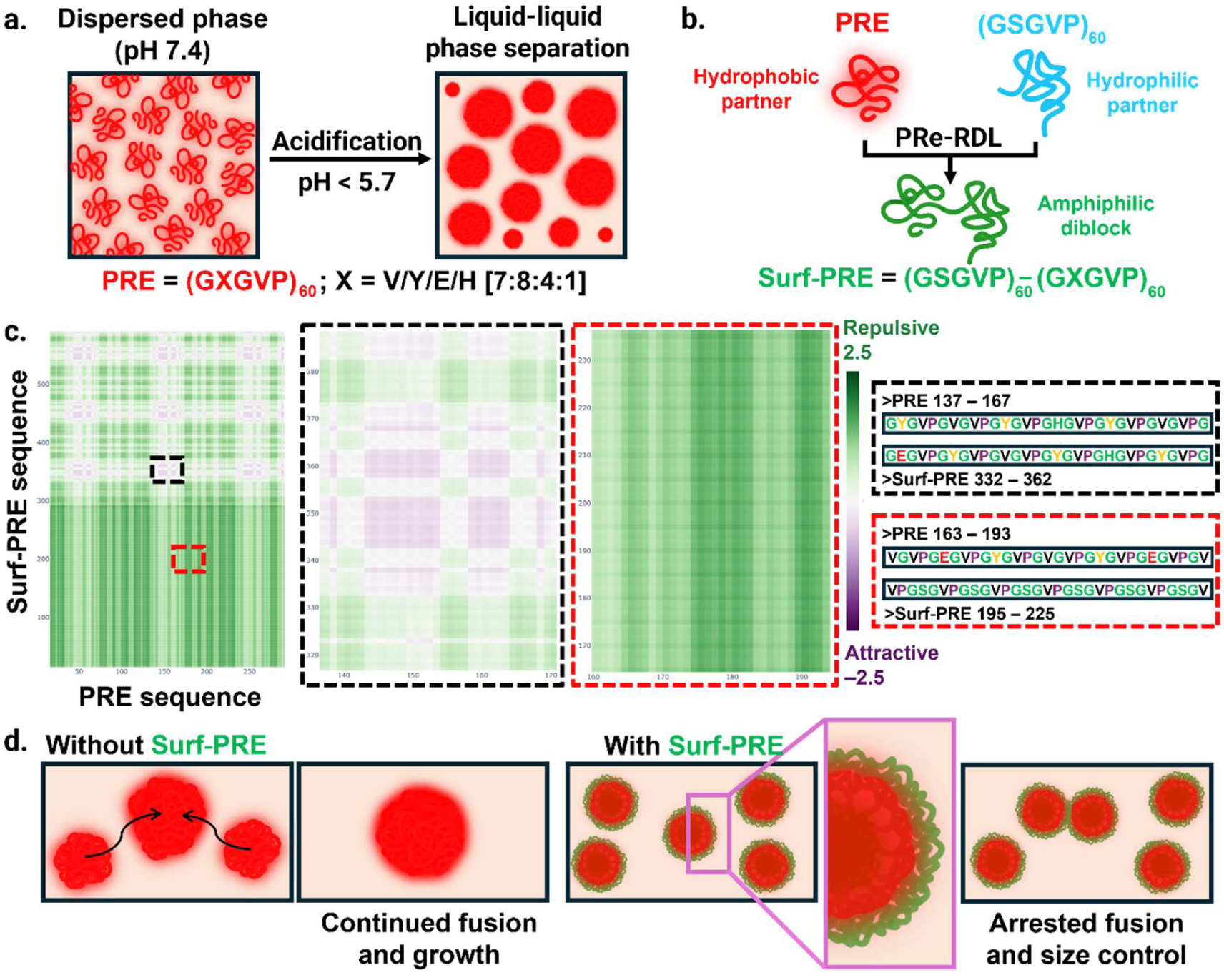
Molecular design and working hypothesis for Surf-PRE-mediated size control of PRE MOs. a) Schematic showing pH-triggered LLPS of PRE. PRE, composed of the repeat sequence (GXGVP)₆₀, remains dispersed at pH 7.4 and undergoes condensation into MOs upon acidification to pH < 5.7. b) Surf-PRE was generated using PRe-RDL, in which the hydrophilic (GSGVP)₆₀ block was genetically fused to PRE, yielding an amphiphilic diblock composed of a PRE-like (GXGVP)₆₀ segment and a (GSGVP)₆₀ segment. c) Pairwise interaction map between Surf-PRE and PRE sequences, used to guide the Surf-PRE design. Green and purple regions indicate repulsive and attractive interaction scores, respectively. The magnified windows highlight the predicted repulsive interaction between the hydrophilic (GSGVP)_60_ block of Surf-PRE and PRE, and the predicted attractive interaction between the PRE-like (GXGVP)_60_ block of Surf-PRE and PRE. d) Proposed model for Surf-PRE-mediated size control. Without Surf-PRE, PRE MOs are expected to undergo continued fusion and growth. With Surf-PRE, the amphiphilic diblock design is hypothesized to modulate the MO boundary and limit coalescence, thereby promoting arrested fusion and size control.

In order to prevent fusion and control the size of MOs through interfacial stabilization, we designed an amphiphilic surfactant composed of protein blocks, called Surf-PRE. The design aimed to balance affinity for PRE condensates with sufficient hydrophilicity to promote interfacial activity. Surf-PRE was designed as PRE-(GSGVP)_60_, where the PRE block provides affinity for the dense phase, while the serine-rich ELP block, (GSGVP)_60_, is hydrophilic and thereby promotes surface activity (Figure 1b, Table S1). The Surf-PRE construct was generated via recursive directional ligation through Recursive Directional Ligation by Plasmid Reconstruction (PRe-RDL)^50^, which allowed the PRE sequence to be genetically fused to the hydrophilic (GSGVP)_60_ block. The resulting plasmid was verified by Sanger sequencing using the primers listed in Table S2. It was subsequently expressed and purified by ITC (see Methods). To evaluate whether this block-level design could generate differential interactions with PRE, we used FINCHES Intermaps^51^ to generate a predicted intermolecular interaction map between Surf-PRE and PRE (Figure 1c). Intermolecular interactions were predicted using FINCHES-online with the Mpipi force field^52,53^. In the Surf-PRE/PRE intermap, the PRE-like block of Surf-PRE showed predicted attractive interactions with PRE, whereas the hydrophilic (GSGVP)_60_ block showed predicted repulsive interactions with PRE. This interaction contrast supported our working hypothesis that Surf-PRE could influence MO growth by modulating interactions at or near the MO boundary, thereby limiting coalescence and promoting size control (Figure 1d). It should be noted that since both PRE and Surf-PRE contain cysteine at their C-terminus, they can form disulfide bonds with each other or with themselves under the non-reducing conditions used throughout these experiments.

To examine the effect of Surf-PRE on MO stabilization, condensation of PRE was performed in the presence of different amounts of Surf-PRE, expressed as *R =* [Surf-PRE]/[PRE], while keeping the PRE concentration constant at 20 µM. PRE condensation was induced using a sharp pH trigger by adding hydrochloric acid (HCl; final concentration 50 mM, corresponding to pH 1.3). Under these conditions, 20 µM PRE rapidly underwent LLPS and formed spherical MOs within minutes. Spinning-disk confocal microscopy (SDCM) images in Figure 2a,b show the resulting condensates after 2 h and 24 h, respectively, with increasing *R* values arranged from left to right. In the absence of any stabilizing additive, the MOs readily fused upon contact, producing large, micrometer-sized condensates with a broad size distribution. This behavior is evident at *R =* 0 in Figure 2a,b, where large PRE-rich condensates were observed at the bottom of the imaging chamber after 2 h and 24 h, respectively. In contrast, the corresponding bulk images shown in the insets contained no visible condensates, indicating that the MOs rapidly sedimented. The *R* values tested further ranged from 0.005 to 0.10, corresponding to 0.1–2 µM Surf-PRE. At the lowest tested *R* value of 0.005, Surf-PRE-stabilized MOs already became significantly smaller compared with PRE-only MOs. Contrary to MOs formed without Surf-PRE, where the condensates were observed mainly at the bottom of the sample chamber, smaller MOs stabilized by Surf-PRE were distributed more homogeneously in the bulk, as seen from the insets in Figure 2a,b. At higher *R* values, this trend became much stronger, with the formation of smaller and more uniformly dispersed Surf-PRE-stabilized MOs that remained suspended within the solution over time. Thus, SDCM imaging provided strong evidence for the stabilization of the MO interface by Surf-PRE molecules, impeding further growth.

**Figure 2.**
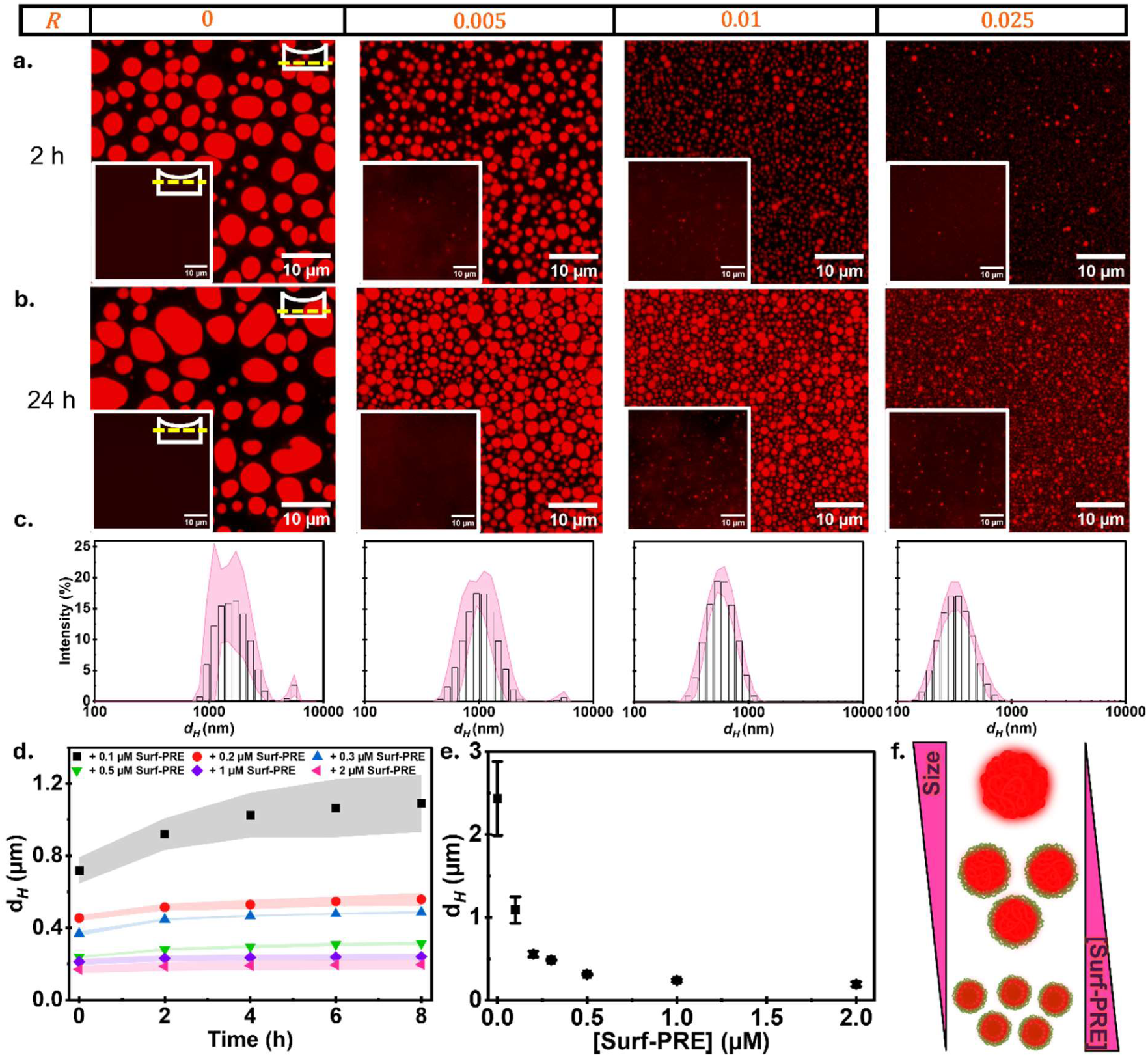
The size of PRE-based MOs is directly regulated by amphiphilic Surf-PRE peptides. a,b) Fluorescence microscopy images of PRE condensates, in the absence or presence of Surf-PRE, 2 h (a) and 24 h (b) after the pH trigger. R denotes the molar concentration ratio, i.e., R=[Surf-PRE]⁄[PRE]. Insets show field-of-views in the bulk of the solution. In the absence of Surf-PRE, MOs readily fuse and grow over time. Increasing concentrations of Surf-PRE result in smaller, highly monodispersed MO populations. c) Size distributions of the condensates, as assessed by DLS, after an 8-hour incubation at different Surf-PRE concentrations. d) Time-dependent evolution of the hydrodynamic diameter of 20 µM PRE condensates in the presence of different Surf-PRE concentrations, showing their stabilization effect. e) Hydrodynamic diameter of MOs as a function of Surf-PRE concentration, 8 h after the pH trigger, highlighting the rapid decrease in the size of the assemblies before reaching a plateau around 200 nm. f) Schematic representation of the Surf-PRE-dependent regulation of MO size. Data are represented as mean ± s.d.; n = 2 independent measurements in each case.

To quantify the condensate size distributions observed by fluorescence microscopy, we employed dynamic light scattering (DLS; see Methods for details). The representative DLS distributions after 8 h are shown in Figure 2c. In the absence of Surf-PRE, the PRE condensates continued to grow, exceeding several micrometers in diameter, consistent with their rapid fusion and sedimentation observed by fluorescence microscopy. In contrast, the addition of Surf-PRE led to strict concentration-dependent size control. At *R =* 0.005, the hydrodynamic diameter (*d_hyd_*) after 2 h was 718 nm with a polydispersity index (PDI) of 0.05. The diameter increased to ≈1100 nm after 8 h, corresponding to a 53% increase in *d_hyd_*, indicating that fusion was reduced but not completely suppressed. At *R =* 0.010, the diameter was reduced to ≈450 nm with a PDI of 0.03, and the subsequent increase over 8 h was limited to 100 nm, showing a stronger stabilization effect. At *R* = 0.025, *d_hyd_* was further suppressed to ≈237 nm with a PDI of 0.04, and increased only by 75 nm after 8 h. This limited increase in *d_hyd_*, together with the narrow size distributions, indicates that higher Surf-PRE concentrations more effectively suppress further coalescence and growth of the MOs. We note that the slight discrepancy between the *d_hyd_* obtained from DLS measurements and the larger condensates observed via fluorescence imaging arises because the latter is biased towards condensates that have settled at the bottom of the visualization chamber; indeed, imaging in the bulk of the solution yields a much more representative sample (see the insets in Figure 2a,b), consistent with DLS measurements.

The evolution of *d_hyd_* at different *R* values over 8 h is shown in Figure 2d, which clearly shows that size arrest was reached rapidly and maintained over several hours. For each Surf-PRE concentration, the MO diameter increased only gradually after the initial formation period, and the extent of this growth markedly decreased as the Surf-PRE concentration increased. This concentration-dependent suppression of growth suggests that Surf-PRE stabilizes the MO interface and limits droplet coalescence, with qualitative and quantitative estimates of interfacial coverage discussed later in the coming sections. At lower Surf-PRE concentrations, fewer Surf-PRE molecules are available during MO formation, allowing more extensive condensate growth and occasional fusion before the interface becomes sufficiently stabilized. At higher concentrations, a greater absolute amount of Surf-PRE can associate with the newly formed MO interface, leading to earlier size arrest and the formation of smaller and more stable MOs. Plotting the hydrodynamic diameter as a function of Surf-PRE concentration further highlights the tunability of the system (Figure 2e). The MO diameter decreased steeply with increasing Surf-PRE concentration and reached a lower limit of *d_hyd_* ≈200 nm at Surf-PRE concentrations above 1 µM. This trend suggests that MO size is governed by the balance between PRE condensation and the amount of Surf-PRE available during droplet formation. Together, these results show that Surf-PRE acts as an efficient protein-based interfacial stabilizer for PRE-rich MOs. By increasing the Surf-PRE concentration, the MO diameter could be continuously reduced from µm-sized condensates to near-200 nm assemblies while maintaining very low PDI values. Thus, Surf-PRE provides a simple and programmable route to tune the size and colloidal stability of PRE-based MOs (Figure 2f).

Given the strong dependence of the LLPS process on pH, we asked whether the acidification rate could also affect MO dynamics and the resulting size control. For this, we used glucono-δ-lactone (GDL), which hydrolyzes gradually in aqueous solution to lower the pH over time. Under the experimental conditions used here, 84.2 mM GDL reduced the pH from 7.4 to 2.8 over the course of 8 h (Supplementary Figure S2). Interestingly, while Surf-PRE retained its ability to control MO size under gradual acidification, the corresponding MO sizes at equivalent R values were consistently larger than those obtained using the sharp acidification trigger (Supplementary Figure S3). The dependence on Surf-PRE concentration was also less pronounced under GDL-triggered acidification. Moreover, once the condensates reached their characteristic size, no substantial further growth was observed, indicating that Surf-PRE arrested MO growth under gradual acidification. Together, these results show that Surf-PRE remains capable of stabilizing MO size under gradual acidification, although the final MO size is larger and less sensitive to Surf-PRE concentration than under sharp acidification. Thus, Surf-PRE-mediated size control depends on two experimentally tunable parameters: the amount of Surf-PRE available for interfacial stabilization and the rate at which PRE condensation is triggered.

### Surf-PRE molecules form a dynamic interface at the MO boundary

While the observed behavior is consistent with surfactant-like stabilization, the molecular basis of Surf-PRE action cannot be inferred from size measurements alone. We therefore examined the localization of Surf-PRE and how its spatial distribution contributed to interfacial stabilization. Surf-PRE was fluorescently labelled with Alexa Fluor 488 (AF488) through cysteine-maleimide conjugation. We examined MOs formed at *R* = 0.005; this condition was chosen because higher Surf-PRE concentrations produced much smaller MOs, making them difficult to image. Confocal laser scanning microscopy (CLSM) revealed a clear enrichment of Surf-PRE at the boundary of PRE MOs (Figure 3a). The AF647-PRE signal was present throughout the dense phase, while the AF488-Surf-PRE signal appeared as a ring-like layer around the condensates. The individual channels in Figure 3b,c further confirmed this spatial distribution, showing a homogeneous PRE signal within the condensates and a strongly enriched Surf-PRE signal at their periphery. We measured the partition coefficient of PREs in the dense phase to be *K*_PRE, C_ = 454.5 ± 8.9 (*n* = 9 condensates; see Methods for details). However, Surf-PRE was not confined exclusively to the interface, as a weaker AF488-Surf-PRE signal was also detected within the PRE-rich interior. This localization pattern was further emphasized by the three-dimensional reconstruction, which showed a Surf-PRE-enriched layer surrounding the MOs and suggested a core-shell-like architecture (Figure 3d). A representative line profile across a single MO confirmed the same trend, with elevated Surf-PRE intensities at the MO boundary compared with the condensate interior (Figure 3e). We quantified this enrichment in terms of partition coefficients of Surf-PRE at the MO interface and within the MO interior (Figure 3f). As expected, Surf-PRE was strongly enriched at the interface, with *K*_Surf-PRE, I_ = 190.3 ± 10.6 (*n* = 9 condensates). Surf-PRE was less strongly, but still notably, enriched within the MOs, with *K*_Surf-PRE, C_ = 164.6 ± 8.7 (*n* = 9 condensates). This localization behavior agrees well with its diblock architecture: the PRE-like segment interacts favorably with the PRE-rich dense phase, while the serine-rich ELP segment has more favorable interactions with the surrounding aqueous phase. Together, these two features allow Surf-PRE to associate with the MO interior while still favoring the interface.

**Figure 3.**
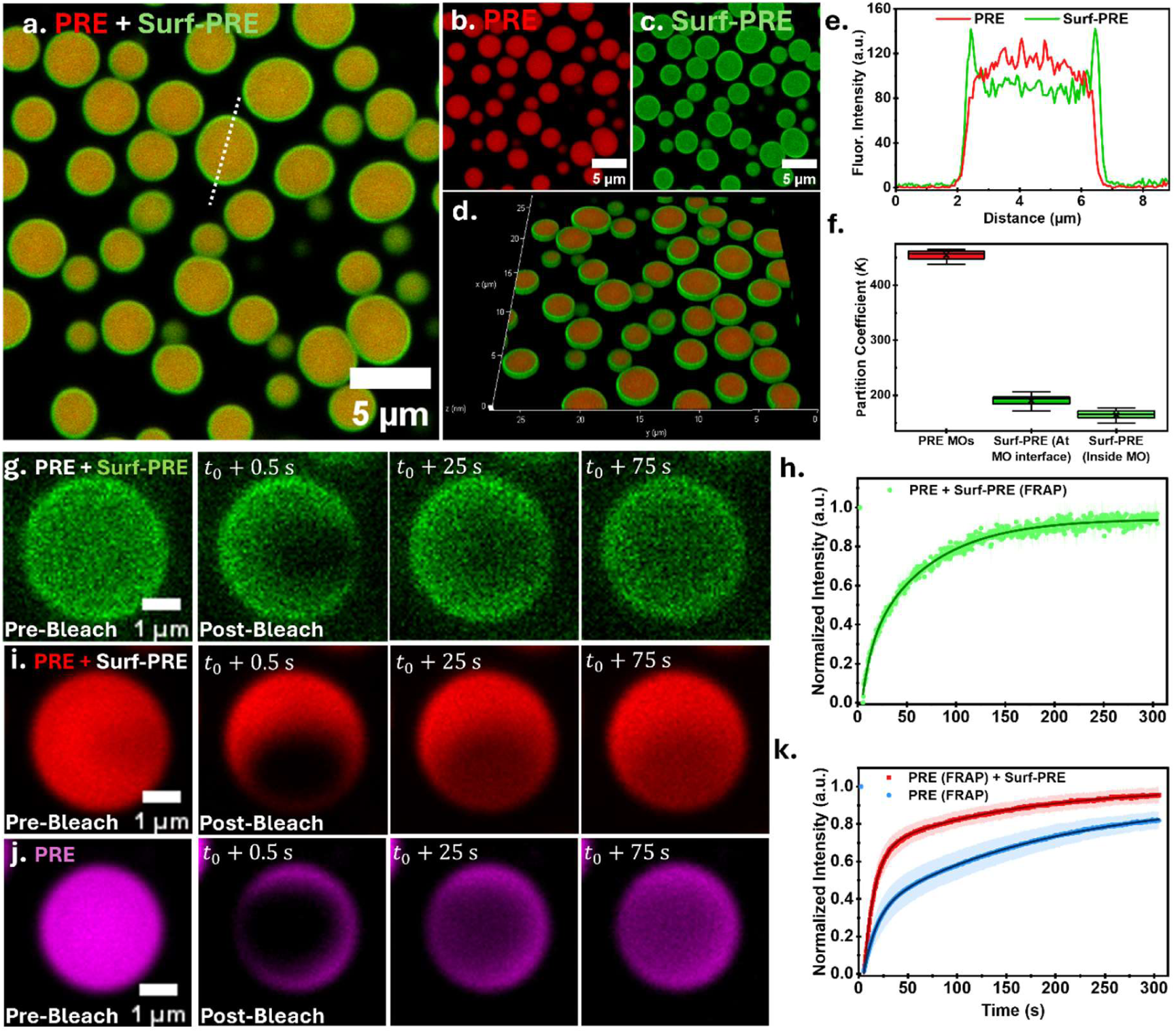
Spatial distribution and fluorescence recovery dynamics of Surf-PRE-stabilized MOs. a) CLSM image of Surf-PRE-stabilized MOs, showing homogeneous distribution of PRE throughout the condensate together with strong Surf-PRE accumulation at the interface. The dashed line indicates the region analyzed in e. b,c) Individual fluorescence channels for PRE (b) and Surf-PRE (c), respectively. d) Single-slice view after 3D rendering of the confocal image, highlighting Surf-PRE enrichment at the MO interfaces. e) Line scan of PRE and Surf-PRE fluorescence intensity across a condensate, showing a homogeneous PRE signal throughout the MO, in contrast to the enriched Surf-PRE signal at the interface. f) Partition coefficients of PRE and Surf-PRE in the indicated regions. From left to right, the plotted values correspond to PRE in the dense phase (red), Surf-PRE at the MO interface (dark green), and Surf-PRE inside the MO (light green), showing preferential accumulation of Surf-PRE at the condensate interface; n = 9 condensates. g,h) FRAP time-lapse images showing pre-bleach and post-bleach events after bleaching Surf-PRE in Surf-PRE-stabilized MOs (g), and the corresponding fluorescence recovery curve (h). i-k) FRAP time-lapse images showing pre-bleach and post-bleach events after bleaching PRE in Surf-PRE-stabilized Mos (i) and PRE in non-stabilized MOs (j), with the corresponding fluorescence recovery curves shown in (k). All recovery curves are fitted with a biexponential function (solid lines). Condensates were prepared with 20 µM total PRE, of which 4 mol% was AF647-PRE, and 0.1 µM AF488-Surf-PRE, corresponding to 0.5 mol% relative to PRE. For FRAP analyses, 3 condensates were analyzed. Data are represented as mean ± s.d. (error bars and shaded areas).

We next asked whether the relative distribution of Surf-PRE between the MO interface and the PRE-rich interior changed with Surf-PRE concentration. We reasoned that, at lower Surf-PRE concentrations, the available molecules might preferentially accumulate at the interface, leading to stronger interfacial enrichment relative to the dense phase. To test this, we compared MOs formed at R = 0.005, 0.0025, and 0.0005. CLSM images obtained at the two lower *R* values showed the same overall localization pattern observed at *R* = 0.005, with Surf-PRE enriched at the MO boundary but also clearly present within the condensate interior (Supplementary Figure S4). Appropriate controls confirmed that AF488-Surf-PRE remained enriched at the interface while also partitioning into the condensate interior, whereas AF647-PRE remained homogeneously distributed without interfacial enrichment (Supplementary Figure S5). We therefore quantified the ratio of the Surf-PRE fluorescence intensity at the interface to that within the dense phase. This ratio remained nearly unchanged across the three conditions, with values of approximately 1.2, 1.2, and 1.1 at *R* = 0.005, 0.0025, and 0.0005, respectively (Supplementary Figure S4). These results indicate that Surf-PRE accumulates within the MOs with only a moderate preference for the interface and that this relative distribution remains largely unchanged over the concentration range examined. Nevertheless, this moderate interfacial preference has a pronounced effect on MO stability and size, as increasing the Surf-PRE concentration allows more interface to be covered and results in the formation of smaller, size-arrested MOs.

To determine the molecular mobility within condensates, we performed fluorescence recovery after photobleaching (FRAP) experiments (Figure 3g-k). We photobleached Surf-PRE-stabilized MOs at *R* = 0.005, using spectrally distinct AF647-PRE and AF488-Surf-PRE molecules to independently monitor their dynamics within the same MO (Figure 3g,i; see Methods for details). Additionally, PRE-only MOs without Surf-PRE were subjected to similar imaging and bleaching conditions (Figure 3j). All three recovery profiles were well described by biexponential fits (Figure 3h,k), consistent with the presence of fast and slow recovery components. In Surf-PRE-stabilized MOs, the AF488-Surf-PRE signal recovered rapidly and reached more than 90% recovery within the experimental window (Figure 3h), with a calculated mobile fraction of 0.94. This high mobile fraction indicates that Surf-PRE remains highly mobile rather than forming an immobile interfacial shell and suggests that the Surf-PRE-enriched boundary is capable of dynamically exchanging with surrounding unbleached regions and/or the dilute phase. The AF647-PRE signal within the same Surf-PRE-stabilized MOs also showed a pronounced initial recovery followed by a slower, gradual increase toward a plateau within the experimental window (Figure 3i,k). The fitted mobile fraction was 0.99, indicating that nearly the entire PRE population remained mobile. This biphasic recovery is consistent with multiple contributions to PRE redistribution, potentially including local diffusion, exchange with the surrounding phase, and rearrangement within the PRE-rich dense network. This interpretation is consistent with PRE acting as the scaffold of the MO phase, where transient intermolecular interactions may slow molecular redistribution. We then carried out FRAP on MOs formed without Surf-PRE, in which AF647-PRE recovered more slowly than in Surf-PRE-stabilized MOs and reached a lower normalized intensity within the experimental window (Figure 3j,k). The fitted mobile fraction also decreased to 0.94. These findings suggest that Surf-PRE not only suppresses MO coalescence but also modifies PRE dynamics within the dense phase.

To more directly test molecular exchange across the interface, whole-droplet FRAP was performed by bleaching the AF647-PRE and AF488-Surf-PRE signals over the entire droplet. In this geometry, fluorescence recovery mainly reports molecular influx from the surrounding dilute phase and exchange with nearby unbleached condensates. Recovery began at the droplet surface and proceeded inward (Supplementary Figure S6). Line intensity profiles further clarified this pattern, with fluorescence appearing first at the periphery of the bleached droplet before becoming more uniform across the droplet interior. Importantly, even in whole-droplet FRAP experiments, PRE-only MOs recovered more slowly, with a noticeable delay before fluorescence appeared throughout the bleached condensates. These results strongly support the notion that Surf-PRE increases molecular exchange between the dilute and MO phases. One possible explanation for the enhanced recovery is that, after the pH trigger, Surf-PRE does not exist only as an interfacial layer on PRE MOs but may also remain present in the surrounding dilute phase as dynamic Surf-PRE-rich nanoscale assemblies. Because of its amphiphilic architecture, Surf-PRE may also remain in the surrounding dilute phase as weak micelle-like assemblies, conceptually reminiscent of the weak micelles reported for ELP diblock copolymers previously^54^. Such dynamic assemblies could act as exchange-competent reservoirs for Surf-PRE and PRE molecules, facilitating their transfer between the dilute phase and the bleached MOs. In this view, Surf-PRE does not form a static coating around the MO; rather, it creates a dynamic boundary that stabilizes the condensate interface while still permitting molecular exchange. Together, confocal imaging and FRAP results show that Surf-PRE performs two coupled functions: enrichment of Surf-PRE at the MO interface provides a stable boundary that prevents coalescence, while the dynamic nature of this boundary allows faster intra-and inter-condensate exchange of PRE molecules. In other words, Surf-PRE physically stabilizes PRE MO boundaries without arresting material exchange.

### Sub-Compartmentalization of synthetic cells using Surf-PRE-coated MOs

After confirming the Surf-PRE-mediated regulation of MO size in bulk, we proceeded to utilize the strategy to sub-compartmentalize synthetic cells. We used double emulsions (DEs) - artificial compartments with a cell-sized aqueous interior (14–17 pL) surrounded by a thin oil shell - as our synthetic vesicles. DEs were produced using a previously established on-chip microfluidic platform^48,55,56^, about 25–30 µm in diameter and encapsulating appropriate amounts of PRE and Surf-PRE into their lumen (Figure 4a; see Methods for details). Importantly, the thin oil shell allows pH equilibration between the lumen and the external medium within minutes, allowing phase separation to be triggered by simply modulating the external pH, while directly monitoring the MO formation inside. To trigger LLPS, the external aqueous phase was changed to an acidic feeding solution (100 mM HCl, 50 mM NaCl, 1% Tween-20). Upon reaching the transition pH of 5.7, typically within 10 mins, coacervation commenced within DEs. Without Surf-PRE, the condensates formed throughout and merged to ultimately form a single MO within each DE. However, when Surf-PRE was present, several distinct MOs remained within each DE even after 24 h, indicating that the same design principle used for bulk MO size control could be successfully replicated in cell-sized micro-confinements.

**Figure 4.**
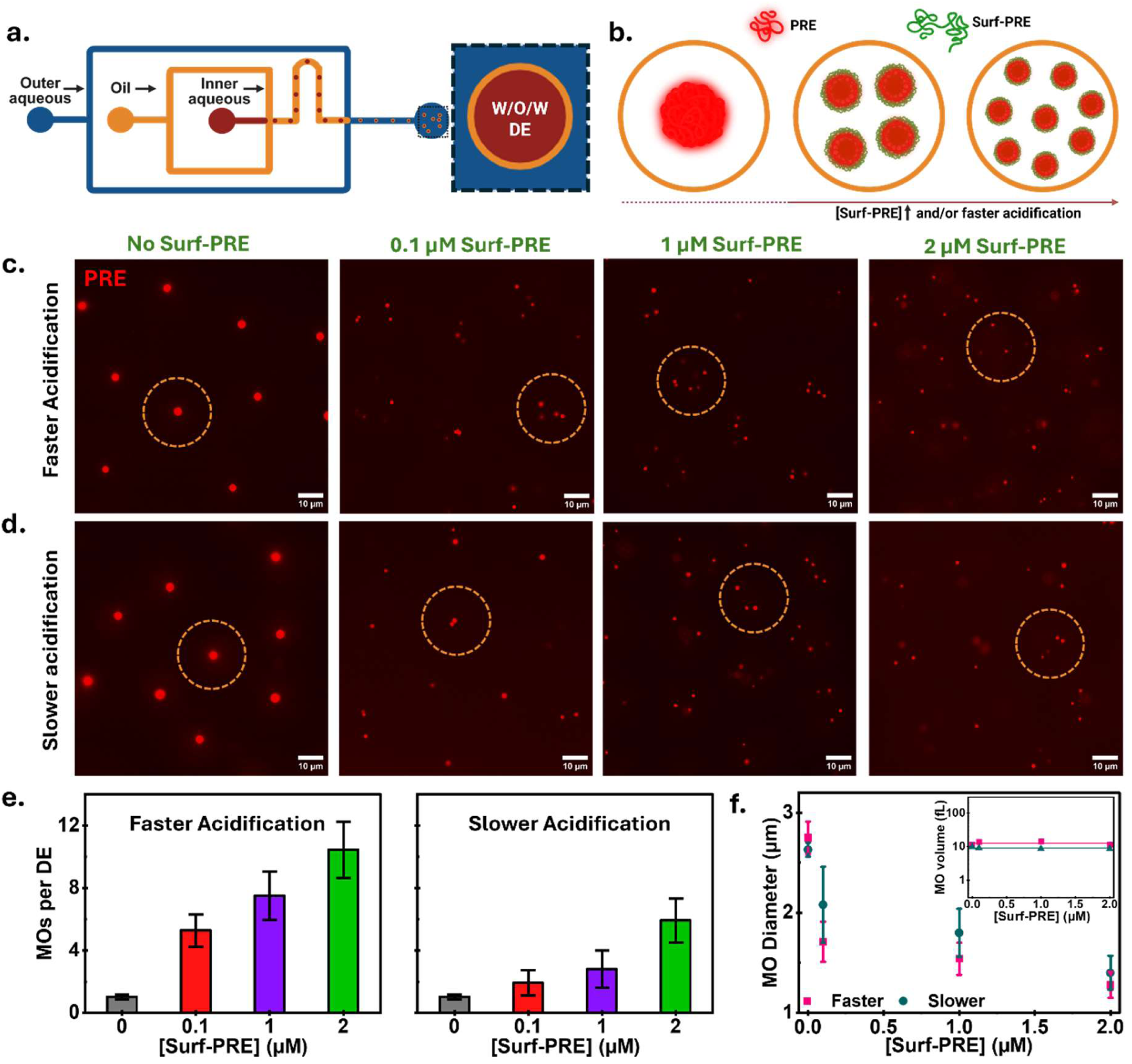
Surf-PRE-mediated subcompartmentalization within synthetic cells. a) Schematic of the microfluidic device used for the generation of water-in-oil-in-water double emulsions (W/O/W DEs). b) Schematic showing pH-triggered MO formation and regulation inside the DEs. Increasing the Surf-PRE concentration and/or acidification rate results in a larger number of smaller MOs. c,d) Representative fluorescence micrographs showing PRE-rich MOs formed inside the DEs at increasing Surf-PRE concentrations under faster (c) and slower (d) acidification conditions. The images were acquired 24 h after the DE production and represent the final state of the system. Dashed circles outline representative DEs. e) Mean number of MOs per DE as a function of Surf-PRE concentration under faster and slower acidification conditions (n = 36 DEs per condition). f) Mean MO diameter as a function of Surf-PRE concentration under faster and slower acidification conditions, showing a decrease in MO size with increasing Surf-PRE concentration. For faster acidification, n = 24 condensates for 0 µM Surf-PRE and n = 100 condensates for the remaining Surf-PRE concentrations were analyzed. For slower acidification, n = 21 condensates for 0 µM Surf-PRE and n = 51 condensates for remaining Surf-PRE concentration were analyzed. The inset shows the corresponding PRE MO volume as a function of Surf-PRE concentration. The inner aqueous phase consisted of 20 µM PRE and the respective Surf-PRE concentrations. The oil phase consisted of a fluorinated oil. For fast acidification, the outer aqueous phase consisted of 100 mM HCl, 50 mM NaCl, and 1% Tween-20, whereas for slow acidification, it consisted of 10 mM HCl, 140 mM NaCl, and 1% Tween-20. Data are presented as mean ± s.d.

To determine the extent of tuning on the internal organization, we quantified the number of MOs formed within different DE populations containing increasing Surf-PRE concentration, 24 h after post-trigger (Figure 4e). Absence of Surf-PRE, as mentioned above, resulted in a single MO in virtually all the DEs. Already at 0.1 µM Surf-PRE, approximately 5 compartments per synthetic cell were observed. Increasing the Surf-PRE concentration to 1 and 2 µM further increased this number to approximately 8 and 10 MOs per DE, respectively. A representative time-lapse acquired at 2 µM Surf-PRE shows the formation and persistence of multiple MOs during acidification (Supplementary Video 1). The DE system also allowed us to estimate the volume fraction of the PRE condensates and, together with the measured partition coefficients, the amount of PRE present in the dense phase versus the dilute phase (see Supplementary Note S1 for details). At 0.1 µM Surf-PRE, the MOs together occupied approximately 0.1% of the DE volume, and considering an estimated 450-fold enrichment of PRE within the condensates, about 29% of the total PRE resided in the dense phase. It should be noted that these Surf-PRE-stabilized MOs were highly stable and showed no sign of coalescence, even when constantly diffusing within the micro-confinement presented by DEs, over at least a 24 h period. Thus, the number of internal membraneless compartments can be programmed by Surf-PRE concentration, giving modular control over the degree of sub-compartmentalization within each synthetic cell.

Similar to bulk experiments, we tested whether the rate of pH trigger also had an impact on MO formation by using a slower acidification trigger (10 mM HCl, 140 mM NaCl, and 1% Tween-20). As with sharp acidification, DEs without Surf-PRE predominantly formed a single PRE-rich MO, confirming that in the absence of interfacial stabilization, a single MO was formed regardless of the severity of the trigger. When Surf-PRE was present, while the number of internal MOs remained tunable, the final MO number remained substantially lower than obtained under rapid acidification (Figure 4e). At 0.1, 1, and 2 µM Surf-PRE, the slower trigger yielded approximately 2, 3, and 5 MOs per DE, respectively. This difference corroborates well with bulk results, consistent with the nucleation-and-growth behavior (Figure 2). Thus, for the same Surf-PRE concentration, the balance between nucleation, growth, and interfacial stabilization could be shifted by modulating the rate of pH change, leading to a different number of final MOs inside the DEs. The different number of MOs formed should also be reflected in their average size. Indeed, a plot of MO diameter as a function of Surf-PRE concentration confirmed this (Figure 4f). The total volume occupied by PRE-rich MOs within each DE (product of the average number of MOs and the average MO volume) showed a steady value regardless of the Surf-PRE concentrations and the acidification rate, consistent with the condensed phase being distributed among progressively more numerous and smaller MOs (Figure 4f, inset).

To conclude, the degree of sub-compartmentalization within synthetic cells was efficiently turned via two knobs: Surf-PRE concentration and the acidification rate, i.e., the trigger kinetics. We now turned our attention to assessing the functionality of these MOs.

### Size-controlled sub-µm MOs enhance enzymatic reaction kinetics

Given the vital role of MOs in providing suitable microenvironments for specific biochemical activities inside cells, we investigated the potential of our size-arrested MOs in supporting enzymatic reactions. We selected acid phosphatase (AcP) as a model enzyme and DiFMUP as a fluorogenic substrate. AcP hydrolyses non-fluorescent DiFMUP to fluorescent DiFMU (Supplementary Figure S7), allowing the reaction to be followed in real time by fluorescence microscopy and spectroscopy (see Methods for details). It should be noted that neither AcP nor DiFMUP was tagged with any PRE-like sequence to enhance their partition inside the MOs. This allowed us to test whether PRE MOs could influence reaction kinetics through passive partitioning and local enrichment of reaction components, rather than by forced recruitment to the phase-separating scaffold. This scenario is closer to many cellular MO systems, where enzymes, substrates, cofactors, or reaction products are often recruited as clients through non-covalent interactions and selective partitioning.

We first confirmed that no fluorescent product was generated in the absence of AcP, establishing that DiFMU formation required enzymatic hydrolysis (Figure 5a). Indeed, upon addition of AcP, fluorescence increased after an hour and became strongly enriched within the PRE-rich condensates (Figure 5a). Quantification showed that the fluorescence signal increased almost eight-fold within the MOs but only three-fold in the surrounding dilute phase (see Methods for details; Figure 5b). The increase in reaction-associated fluorescence within MOs supports their role as localized reaction hubs, likely through the recruitment of AcP and/or DiFMUP and the promotion of enzymatic hydrolysis within the dense phase or at the dense-dilute interface.

**Figure 5.**
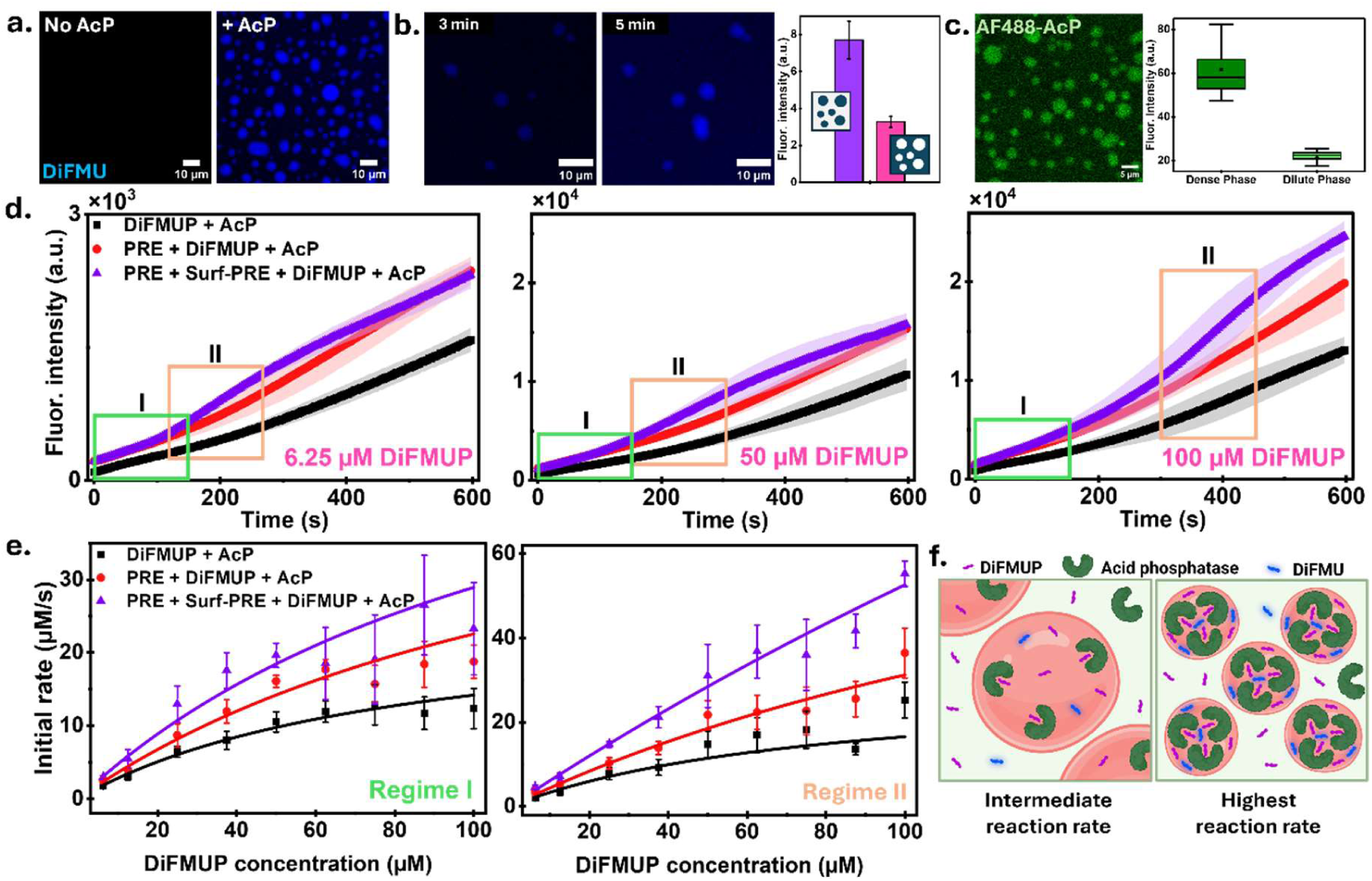
Size-controlled sub-µm MOs act as reaction hubs and show enhanced catalytic activity for acid phosphatase. a) Representative multiphoton fluorescence images of PRE MOs without and with AcP, showing product formation only in the presence of AcP. The key feature is the appearance of DiFMU fluorescence within PRE MOs after AcP addition, indicating MO-localized DiFMUP hydrolysis. b) Time-lapse showing DiFMU formation as a result of DiFMUP hydrolysis by AcP. Time stamps indicate the time elapsed after AcP was added to the DiFMUP-containing MO population. Bar graph quantifying the DiFMU fluorescence in the condensates (n = 5 condensates) versus the dilute phase. c) CLSM micrograph of AF488-labeled AcP showing its localization within PRE MOs, together with quantification of AcP fluorescence intensity in the dense and dilute phases (n = 52 MOs). d) Kinetics of DiFMUP hydrolysis in a homogeneous solution without MOs (black), non-arrested MOs formed from 25 µM PRE (red), and size-arrested MOs containing 1 µM Surf-PRE (purple), at DiFMUP concentrations of 6.25, 50, and 100 µM. Fluorescence measurements were initiated immediately after AcP addition. Regimes I and II indicate the time windows used for the reaction-rate analyses shown in e (n = 5 independent bulk reactions for each condition). e) Reaction rates calculated from regime I (left) and regime II (right) as a function of DiFMUP concentration. Solid lines indicate Michaelis-Menten fits (R² > 0.95). f) Schematic illustrating enhanced AcP-catalyzed DiFMUP hydrolysis within MOs and the further increase in reaction rate in smaller, size-arrested MOs containing Surf-PRE. Data are presented as mean ± s.d., represented by error bars or shaded areas.

To visualize AcP localization, the enzyme was labelled with Alexa Fluor 488 C5 maleimide at a 1:1 dye-to-protein molar ratio to favour predominantly single-site labelling (see Methods for details). We then incubated 1 µM AF488-AcP with 25 µM PRE and examined its localization by CLSM. AF488-AcP fluorescence was ≈2.8-fold higher inside PRE MOs than in the surrounding dilute phase, indicating that the enzyme preferentially partitions into the condensate phase (Figure 5c). Upon confirming that AcP localizes within MOs and can drive DiFMUP hydrolysis within the condensates, we proceeded to measure the enzymatic activity over a wide range of DiFMUP concentrations (6.25–100 µM) using a microplate reader (see Methods for details). The concentrations of PRE, Surf-PRE, and AcP were kept constant at 25 µM, 1 µM, and 0.05 U, respectively. Three reaction environments were compared: bulk solution (DiFMUP and AcP), MOs without size control (PRE, DiFMUP, and AcP), and size-arrested MOs (PRE + 1 µM Surf-PRE, leading to ∼250-nm-sized condensates, DiFMUP, and AcP). Control samples prepared without AcP showed no detectable increase in fluorescence over the same measurement period, confirming that the observed signal arose from enzyme-catalyzed DiFMUP hydrolysis (Supplementary Figure S8). Notably, across all substrate concentrations tested, DiFMU formation proceeded fastest in samples containing size-arrested MOs, followed by MOs without size control and finally the bulk solution (Figure 5d; Supplementary Figure S9). The enhanced reaction rate in the presence of condensates clearly indicated that the MOs provided local enrichment of reaction components, thereby increasing the effective concentration of the reacting species and shifting the apparent reaction kinetics away from those observed in a homogeneous bulk solution. To quantify the reaction kinetics, we used the initial regime of 150 s and fitted the reaction rate-versus-substrate plots using the Michaelis-Menten model (Figure 5e, see Methods for details). While the apparent reaction rate increased with DiFMUP concentration for all three reaction conditions, the magnitude of the rate enhancement depended strongly on the reaction environment. As can be seen from Supplementary Table S3, the apparent maximum velocity (*V_max_*) increased from 26 µM s⁻¹ in bulk solution to 55 µM s⁻¹ in non-arrested MOs and 67 µM s⁻¹ in size-arrested MOs. Thus, phase separation enhanced the apparent catalytic output, with size-arrested MOs giving the largest increase. While the apparent *K_m_* values were higher in MO-containing samples (>130 µM) compared to bulk (84 µM), the apparent catalytic efficiency, estimated as *V_max_*/*K_m_*, increased from 0.31 in bulk to 0.38 in PRE MOs and 0.50 in Surf-PRE-stabilized MOs, respectively, representing a 23% and 67% increase. This increase indicates that the gain in maximal reaction rate outweighed the accompanying increase in apparent *K_m_*, resulting in an overall enhancement of enzymatic performance within the MOs.

This clear increase in the reaction efficiency in size-arrested MOs merits extra attention, as it suggests that compartment size also contributes to the catalytic output. In the presence of Surf-PRE, PRE MOs are much smaller and more numerous, increasing the total interfacial area between the dense and dilute phases. The resulting higher surface-to-volume ratio may promote faster substrate exchange per unit condensate volume, while shorter intraphase diffusion distances may allow incoming DiFMUP molecules to encounter AcP more efficiently before returning to the surrounding phase. Together, these effects may facilitate substrate replenishment and product release across the MO interface.

Beyond the initial reaction regime, the kinetic traces revealed a second, transient regime in which the effect of MO size became particularly pronounced. Starting at approximately 200 s, the fluorescence signal from size-arrested MOs increased more steeply than that from non-arrested MOs for a period of ∼400 s, before the two MO-containing conditions gradually approached similar fluorescence values at later time points (Figure 5d; Supplementary Figure S9). This transient divergence and subsequent convergence of the kinetic traces produced an “eye-like” regime (addressed as Panel II in Figure 5d), which was consistently observed across the substrate concentrations tested. To quantify this behavior, we extracted apparent reaction rates from the first 150 s of this eye-like regime. The rate enhancement was particularly pronounced in this time window, with Surf-PRE-stabilized MOs clearly separating from both PRE-only MOs and the bulk reaction. Consistent with this observation, the effect of MO confinement was strongest in regime II (Figure 5e). The apparent *V_max_* increased only modestly in bulk solution, from 26 to 30 µM s⁻¹, but rose to 100 µM s⁻¹ in PRE MOs and 352 µM s⁻¹ in Surf-PRE-stabilized MOs. The corresponding apparent *K_m_* values also increased, reaching 220 µM for PRE MOs and 570 µM for Surf-PRE-stabilized MOs, compared with 81 µM in bulk solution. A similar increase in apparent *K_m_* was also observed in the initial regime, indicating that this deviation is not unique to the eye-like window but is a general feature of AcP catalysis within MOs. Instead, the fitted parameters likely report a combined contribution from enzyme turnover, substrate partitioning, product accumulation, and transport between the dense and dilute phases. Notably, although the fit quality for the homogeneous reaction decreased in this later time window (R² = 0.94), the PRE-only MO and Surf-PRE-stabilized MO datasets remained well described by the Michaelis-Menten-type fits (R² = 0.98 and 0.99, respectively). This may suggest that the MO phase imposes a more defined local reaction environment, where substrate partitioning and enzyme localization partially maintain an apparent enzyme-saturation relationship even when the assumptions of bulk Michaelis-Menten kinetics are no longer fully satisfied. The apparent catalytic efficiency also remained highest in the Surf-PRE-stabilized MOs, reaching 0.62 compared with 0.45 for PRE MOs and 0.37 for the bulk solution.

Together, these results suggest that the eye-like regime represents a period in which reaction, partitioning, and molecular exchange between the dense and dilute phases are most strongly coupled. In other words, the advantage of smaller MOs is likely present from the beginning but becomes most apparent once substrate redistribution and product formation have established concentration gradients across the MO interface. Under these conditions, the higher surface-to-volume ratio and shorter intraphase diffusion length of size-arrested MOs may support faster substrate replenishment and product release per unit condensate volume than larger PRE-only MOs. The stronger separation observed at higher DiFMUP concentrations is consistent with this interpretation, as a larger substrate reservoir in the dilute phase may sustain these gradients for longer and make the size-dependent difference in MO-mediated turnover more visible. Thus, Surf-PRE-stabilized sub-µm MOs not only enhance the initial apparent reaction rate but also support a transient period of accelerated product formation before the kinetic traces converge at later times.

Overall, the kinetic analysis showed a clear hierarchy in the enzymatic activity. Surf-PRE-stabilized MOs showed the highest reaction rates, followed by µm-sized PRE MOs (Figure 5f), while the bulk reaction was slowest. These results therefore suggest that arresting MO fusion and maintaining smaller condensates enhances enzymatic output more effectively than allowing PRE MOs to grow into larger condensates. More broadly, the data show that compartment size and stability directly influence catalytic performance, with small Surf-PRE-stabilized MOs outperforming both bigger MOs and the homogeneous bulk reaction.

## DISCUSSION

This work describes the design and implementation of a pH-triggered LLPS system that enables exquisite control over the size, and thus the number, of peptide-based condensates formed within a finite reaction volume. Two important consequences of this strategy are shown: facile multi-compartmentalization within synthetic cells by forming a specific number of membraneless organelles, and enhanced enzymatic reaction rates by providing reaction hubs with large interface-to-volume ratios.

LLPS of IDPs (PRE in our case) alone has no control over the MO population, as the formed condensates continue to merge and grow until equilibrium is reached. However, with the addition of an amphiphilic peptide, Surf-PRE in our case, a stable, monodispersed MO population was obtained. This behavior is consistent with the surfactant-like role of Surf-PRE, where its partitioning into the PRE-rich dense phase is accompanied by stronger enrichment at the dense-dilute phase boundary, which can reduce the interfacial free-energy penalty and limit condensate fusion in line with general surfactant adsorption principles^57–59^. Consistent with this interpretation, chemically distinct amphiphilic diblock ELPs and interface-localizing peptides have also been shown to suppress coarsening or alter condensate size distributions^60,61^. Furthermore, increasing the Surf-PRE concentration decreases the MO size, as more Surf-PRE becomes available to stabilize the interface and inhibit fusion, arresting condensates at smaller sizes.

We further show that gradual acidification leads to larger MOs compared to sharp acidification. This observation agrees with previous work showing that condensate nucleation kinetics depend on quench depth, that is, how far a system is driven into the two-phase regime. In this framework, a shallow quench, corresponding to a weaker effective supersaturation close to the phase boundary, leads to slower nucleation and fewer mesoscopic assemblies compared to a deeper quench^62,63^. Applying this framework to our system, sharp acidification may produce a more abrupt increase in supersaturation as PRE crosses the phase boundary, favouring a larger number of nucleation events. Surf-PRE could then stabilize the newly formed interfaces early in the LLPS process, distributing the available PRE among a larger number of MOs and thereby limiting their final size. In contrast, gradual acidification may keep the system close to the transition boundary for longer, allowing fewer MOs to nucleate initially and each MO to grow further before Surf-PRE enrichment becomes sufficient to arrest continued growth. This interpretation could also account for the weaker dependence of MO size on Surf-PRE concentration under gradual acidification, where the nucleation and growth pathway may exert a stronger influence on the final MO size than Surf-PRE concentration alone. However, nucleation rates, transient supersaturation, and Surf-PRE accumulation during MO formation were not directly measured in this study. The proposed mechanism therefore remains a possible explanation for the observed dependence of MO size on the acidification rate rather than a directly demonstrated pathway.

Recent studies have shown that amphiphilic block polymers can spontaneously self-assemble at condensate interfaces and form a protective layer that prevents fusion and coarsening without disrupting the liquid-state properties, selectivity, or biological activity of the condensates^64,65^. Such observations highlight that stabilization of condensate interfaces does not necessarily result in solidification or biological inactivity of the condensates. In our current work, we take advantage of this general principle in a simpler polypeptide-based system, where Surf-PRE is designed to partition into the PRE condensates. Our FRAP data indicate that both PRE and Surf-PRE recover rapidly upon photobleaching, suggesting that Surf-PRE enrichment at the interface does not immobilize the condensate. Instead, they maintain a dynamic molecular exchange with the dense phase and its surroundings. Furthermore, the recovery curves showed a strong fit to a biexponential function, suggesting that the diffusion of PRE and Surf-PRE in the molecularly organized compartments may involve more than one recovery component. Such biexponential kinetics are consistent with previous findings and are usually interpreted as a characteristic of different mobile fractions or a time-dependent change in material properties^66,67^. Finally, the whole-MO photobleaching experiments confirm that the recovery rate of PRE from the dilute state is enhanced in the presence of Surf-PRE.

To demonstrate sub-compartmentalization, we used on-chip produced W/O/W DEs as synthetic cells to encapsulate PREs and Surf-PREs and form MOs inside the DEs by modulating the pH of the external environment, which increases hydronium ion activity in the lumen and lowers the lumen pH below the transition pH, thus inducing coacervation. More broadly, microfluidically generated cell-sized confinements have enabled coacervate formation to be triggered and monitored through controlled transport across the compartment boundary^68,69^. Changing the concentration of Surf-PRE allowed us to control the average number of MOs formed inside each DE, showing a simple yet effective strategy to design multi-compartmentalized synthetic cells. Similar to bulk measurements, we found the rate of acidification to have a strong effect on MO formation: slower acidification led to the formation of larger but fewer MOs compared to fast acidification. Thus, we have two independent parameters to control the MO number and size: Surf-PRE concentration and rate of acidification. A related dependence on the rate of triggering has also been observed in microfluidically generated droplets, where faster cooling resulted in a greater number of condensates forming within the confined volumes^70^.

The bio-functionality of our system is shown through the drastic effect size-controlled condensates have on enzymatic reactions, through the example of the hydrolysis reaction by acid phosphatase. In general, confinement can affect catalysis through mechanisms such as substrate diffusion, local substrate concentration, enzyme stability, and the relation between reaction volume and surface area^71^. However, these effects can vary greatly depending on the specific enzyme, substrate, and composition of the compartment. Of particular interest here are condensates, which can serve as active reaction environments due to their dense internal composition. For example, kinase condensates have been shown to accelerate autophosphorylation through local kinase concentration^72^. In coacervate-based protocells with stabilized compartments, Glucose oxidase (GOx) and Horseradish peroxidase (HRP) loaded into separate condensates supported a cascade reaction^73^, while osmotically triggered organelles formed within lipid vesicles incorporated an ELP–HRP conjugate and accelerated its enzymatic activity^74^. Similarly, microfluidic coacervate organelles containing formate dehydrogenase showed increased reaction rates through enzyme/cofactor partitioning and product exchange with the surrounding aqueous phase^75^. Here, we show that the acid phosphatase reaction rate increases dramatically in the presence of MOs and further when the MO size is restricted. This is consistent with the idea that the dense phase acts as a confined reaction medium, significantly elevating the local enzyme concentration. As a result, the MOs can effectively increase the interaction between the enzyme and substrate molecules, producing a faster reaction than in bulk. Interestingly, the effect is even stronger with size-controlled sub-µm MOs. Since enzyme partitioning is independent of the MO size, this effect results purely from the smaller MO size. A likely explanation is linked to their higher surface-to-volume ratio, creating steep substrate gradients at the MO interface, allowing higher flux of substrate conversion. Release of the product into the surrounding dilute phase, preventing product inhibition, might also aid faster reaction rates.

Overall, our findings suggest that such an ELP-based condensate-surfactant system is not only effective for building multi-compartmentalized artificial cells but also for controlling biomolecular exchange and enhancing enzymatic reactions. Related studies using engineered sticker-spacer polypeptides have shown that condensate viscoelasticity can be tuned through peptide sequence, polymer architecture, and mixture composition, supporting the potential of IDP-based condensates as tunable soft biomaterials^76,77^. Various uses of ELP-based materials in drug delivery systems have been explored^78–80^, suggesting the potential applicability of the presented system as a pH-responsive biomaterial platform, particularly for cargo loading and release. Indeed, pH-responsive drug delivery systems have been widely studied because different biological environments show pH values that differ from normal physiological conditions, including tumour-like microenvironments, endosomes, inflamed tissues, and wounds^81–84^. Furthermore, coacervate-based systems have been studied as materials that can encapsulate and retain different types of molecules, including proteins, nucleic acids, growth factors, small molecules, and therapeutic cargo, with their exchange and release behaviour tuned via environmental triggers such as pH and temperature^85–87^. Using this platform, it may be possible to design pH-responsive depots, sequestration materials, enzyme-loaded condensates, reaction vessels, or artificial-organelle-like compartments within synthetic cells. Moreover, this system could be extended beyond cargo delivery to serve as a bioactive-material device for organizing biochemical reactions. Since condensates are increasingly being explored as reaction compartments, the PRE-Surf-PRE system could contribute to this field by providing a biomaterial platform in which the physical state of compartments, enzyme and substrate enrichment, and molecular exchange can be controlled^88,89^.

## MATERIALS AND METHODS

Polyvinyl alcohol (PVA; 87-90% hydrolyzed, average MW 30-70 kDa), propylene glycol methyl ether acetate (photoresist developer), D-(+)-Glucono-delta-lactone (GDL) and acid phosphatase (from potato) were purchased from Sigma-Aldrich. Phosphate-Buffered Saline (PBS) Tablets (Gibco), Alexa Fluor™ 488 C_5_ Maleimide, Alexa Fluor™ 647 C_2_ Maleimide and EnzChek® Phosphatase Assay Kit were purchased from Thermo Fisher Scientific. Sylgard™ 184 silicone elastomer and curing agent were purchased from Dow. Silicon wafer was bought from Silicon Materials. Photoresist (EpoCore 10) was purchased from Microresist Technology GmbH. Microfluidic accessories, including Tygon® tubing 1/16” ODx 0.02”, biopsy punchers, XXS microfluidic reservoir, and HFE 7500 fluorinated oil containing 2% FluoSurf-C surfactant were purchased from Darwin Microfluidics. Elveflow pressure controller OB1-MK3 was used to control the fluid flow.

### Plasmid Construction

The expression plasmid encoding PRE-*h*-36, referred to here as PRE, was generated as described previously^47^. Surf-PRE was assembled by recursive directional ligation by plasmid reconstruction (PRe-RDL). Briefly, two sequential PRe-RDL cycles were used to expand the ELP block from (GSGVP)_20_ to (GSGVP)_60_. The resulting (GSGVP)_60_ block was then inserted in the N-terminus of the PRE-coding sequence in a subsequent PRe-RDL step, yielding the final Surf-PRE expression construct. All plasmids were verified by Sanger sequencing across the complete coding region before transformation into T7 Express *E. coli* cells (New England Biolabs) by heat shock.

### Recombinant protein expression and purification

PRE and Surf-PRE were expressed in T7 Express *E. coli* cells. A single colony was used to inoculate 25 mL Terrific Broth supplemented with 50 mg L^-1^ kanamycin, and the starter culture was grown overnight at 37 °C with shaking. The overnight culture was used to inoculate 1 L LB medium containing 10 g L^-1^ tryptone, 10 g L^-1^ NaCl, and 5 g L^-1^ yeast extract, supplemented with 50 mg L^-1^ kanamycin. Cells were grown at 37 °C in 2 L baffled Erlenmeyer flasks until the culture reached an OD600 of 0.6–0.8. Protein expression was induced with 1 mM isopropyl β-D-1-thiogalactopyranoside (IPTG), after which the cultures were incubated overnight at 18 °C with shaking. Cells were harvested by centrifugation at 6000 rpm, and the resulting pellet was resuspended in 30 mL ice-cold PBS, pH 7.4, supplemented with 1 mM phenylmethylsulfonyl fluoride (PMSF). Cells were lysed on ice by sonication for 7 min using a 2 s pulse cycle at 80% amplitude (Qsonica Q125, CL-18 probe). The lysate was clarified by centrifugation at 12,500 rpm for 30 min at 4 °C, and the soluble fraction was purified by inverse transition cycling (ITC).

Before ITC, a bake-out step was included to reduce soluble impurities. The clarified lysate was incubated at 40 °C for 15 min, cooled at 4 °C for 30 min to resolubilize the PRE-based proteins, and centrifuged at 12,500 rpm for 15 min at 4 °C. The supernatant was then subjected to three rounds of ITC. For each cycle, NaCl was added to a final concentration of 3 M to trigger phase separation, and the sample was incubated at 40 °C for 30 min with shaking at a 45° angle. The phase-separated protein-rich fraction was collected by centrifugation at 12,500 rpm for 15 min at 40 °C. The supernatant was discarded, and the pellet was resuspended in ice-cold 1X PBS. The sample was then incubated at 5 °C for 30 min with shaking to resolubilize the protein, followed by centrifugation at 12,500 rpm for 15 min at 4 °C to remove insoluble material. The supernatant from this cold spin contained the purified PRE-based protein and was carried forward to the next ITC cycle. The resuspension volume was reduced after each cycle according to the size of the recovered pellet. After three ITC cycles, SDS-PAGE analysis indicated >95% protein purity.

### Matrix-assisted laser desorption/ionization time-of-flight mass spectrometry

Before analysis, protein samples were desalted by serial dilution in Milli-Q water and adjusted to a final concentration of 1 mg mL^-1^. The matrix solution was prepared by dissolving 5 mg of 2,5-dihydroxybenzoic acid (DHB) in 200 µL of a Milli-Q water/acetonitrile mixture containing 0.1% formic acid in the acetonitrile phase (2:1, v/v). For sample spotting, 1 µL of matrix solution was first applied to a ground-steel MALDI target plate (MTP 384 target plate ground steel TF, Bruker), followed by 1 µL of protein sample. The spots were mixed on the plate and dried under a gentle stream of warm air before measurement. Mass spectra were acquired using an ultrafleXtreme MALDI-TOF mass spectrometer (Bruker Daltonics), and the data were analyzed with FlexAnalysis software version 3.4 (Bruker).

### Bioconjugation of AF488-Surf-PRE, AF647-PRE and AF488-AcP

Alexa Fluor 488 C5 maleimide and Alexa Fluor 647 C5 maleimide were dissolved separately in dimethyl sulfoxide (DMSO) to prepare 10 mM dye stocks. PRE, Surf-PRE, and AcP were diluted to approximately 1.5 mg mL^-1^ in 1X PBS, pH 7.4. To reduce accessible cysteine residues and any disulfide-linked species, tris(2-carboxyethyl)phosphine (TCEP) was added at a 10-fold molar excess relative to protein, and the samples were incubated for 1 h at room temperature. Alexa Fluor 488 maleimide was then added dropwise to Surf-PRE at a 10-fold dye-to-protein molar excess, whereas Alexa Fluor 647 maleimide was added to PRE at the same molar excess. For AcP (Sigma-Aldrich), Alexa Fluor 488 maleimide was added at a 1:1 dye-to-protein molar ratio to favor predominantly single-site labeling in view of the potential presence of multiple cysteine residues in the commercially obtained enzyme. The reaction mixtures were protected from light with aluminium foil and incubated overnight at 4 °C with gentle rocking. Unreacted dye was removed by repeated dilution with PBS followed by concentration using 10 kDa molecular weight cut-off centrifugal filters (Amicon). Washing was continued until no detectable dye absorbance was observed in the filtrate by UV-vis spectroscopy. The purified conjugates are referred to as AF488-Surf-PRE, AF488-AcP, and AF647-PRE. Labeled proteins were stored for further use at –20 °C.

### Dynamic light scattering

The hydrodynamic diameter of PRE condensates was measured by DLS using a Malvern Zetasizer Nano series instrument equipped with a 632.8 nm He-Ne laser. Samples were prepared in PBS prepared from Gibco PBS tablets and loaded into a low-volume quartz batch cuvette (ZEN2112), taking care to avoid visible bubbles. Measurements were performed at 20.0 °C after an equilibration time of 30 s. For data acquisition and analysis, the material was defined as protein, with a refractive index of 1.450 and an absorption coefficient of 0.001. The dispersant viscosity was used as the sample viscosity. Scattered light was detected at 173° using the non-invasive backscatter configuration. For each time-point and experiment condition, three measurements were recorded with a 60 s delay between measurements. The measurement position was selected using the seek-for-optimum-position setting, and automatic attenuation was used unless manual adjustment was required to maintain a suitable mean count rate. When adjusted manually, the attenuation was selected to keep the mean count rate within approximately 100–400 kilo counts per second (kcps). For monodisperse samples, the *z*-average diameter obtained from cumulant analysis was used as the representative hydrodynamic diameter. Size distributions are reported as intensity-weighted distributions.

### pH Calibration of GDL Hydrolysis

Gradual acidification by GDL hydrolysis was characterized by monitoring the pH of a GDL solution over 8 h. 750 mg of GDL was dissolved directly in 50 mL of 1x PBS (at pH 7.4), yielding a final GDL concentration of 15 mg/mL (84.2 mM). The solution was continuously stirred at 20 °C, and its pH was measured using a calibrated pH probe at 5 s intervals. Measurements were performed in three independent replicates.

### Fluorescence microscopy of PRE condensates

PRE condensates in bulk solution and within double emulsions (DEs) were imaged using a Nikon Ti2-Eclipse spinning-disk confocal microscope equipped with a pE-300ultra illumination system, a Nikon Plan Apo λ 100x oil-immersion objective, and appropriate Semrock filter sets. Images were recorded with a Prime BSI Express sCMOS camera. For bulk imaging of AF647-labeled PRE condensates shown in Figure 1, fluorescence images were acquired using 10%

LED intensity and a 10 ms exposure time. The same samples were also imaged in bright-field using a 10 ms exposure time and 75% transmitted-light intensity. For DE experiments, bright-field images were acquired with a 10 ms exposure time and 75% transmitted-light intensity. Fluorescence images of condensates inside DEs were acquired using 50% LED intensity and a 50 ms exposure time. Following acquisition, display ranges were adjusted independently for the main panels and corresponding insets to improve visualization

### CLSM imaging of Surf-PRE partitioning

Partitioning of AF488-Surf-PRE at the interface and within AF647-PRE condensates was imaged on a Leica Stellaris STED microscope operated in confocal mode. STED depletion was not used for these measurements. Images were acquired in xyz mode using an HC PL APO CS2 86x/1.20 water-immersion objective, with a zoom factor of 5 and a pinhole setting of 152.1 µm, corresponding to approximately 1 Airy unit. The scan was performed unidirectionally at 400 Hz, with no line or frame averaging. Image stacks were acquired with a 512 x 512 pixel format over a field of view of 27.03 x 27.03 µm, giving a pixel size of 0.053 µm and the *z*-stacks were acquired with a step size of 0.225 µm. Fluorescence was detected sequentially to reduce spectral overlap between the AF488-Surf-PRE and AF647-PRE channels. AF488-Surf-PRE was excited using the 492 nm laser line at 22% intensity and detected with a HyD X detector over 498-582 nm in counting mode. AF647-PRE was excited using the 637 nm laser line at 0.07% intensity and detected with a HyD S detector over 642-806 nm in counting mode. For the single-color control experiments, z-stacks were acquired with a step size of 0.356 µm and two line accumulations. The AF488-Surf-PRE-only control was excited at 491 nm using 10% laser intensity and detected with a HyD S detector over 496–802 nm, whereas the AF647-PRE-only control was excited at 653 nm using 0.10% laser intensity and detected with a HyD S detector over 658–829 nm; both channels were recorded in counting mode. Images were acquired using LAS X software.

### Fluorescence recovery after photobleaching

FRAP experiments were performed on a Leica SP8 multiphoton microscope operated in confocal mode. Multiphoton excitation was not used for these measurements. FRAP was performed separately for AF488-Surf-PRE and AF647-PRE to quantify the recovery of Surf-PRE and PRE, respectively, within PRE condensates. Images were acquired in xyt mode using an HC PL IRAPO 40x/1.10 water-immersion objective, with a zoom factor of 15 and a pinhole setting of 77.2 µm, corresponding to approximately 1 Airy unit. The scan was performed unidirectionally at 400 Hz, with one frame average and two line averages. FRAP movies were acquired with a 304 x 95 pixel format over a 19.38 x 6.01 µm field of view, corresponding to a pixel size of 0.064 µm. Each acquisition contained 600 frames recorded over approximately 300 s, with a frame interval of 0.5 s. For each condensate, an elliptical bleach region of interest (ROI), measuring approximately 1.5 µm along the major axis and 1.0 µm along the minor axis, was selected within the condensate. For AF488-Surf-PRE FRAP, fluorescence was monitored using the 488 nm laser line at 2.0% intensity and detected with an internal PMT over 499-560 nm. Photobleaching was performed within the selected ROI using the 488 nm laser line at 80% intensity. For AF647-PRE FRAP, fluorescence was monitored using the 638 nm laser line at 0.05% intensity and detected with an internal PMT over 650-741 nm. Photobleaching was performed within the selected ROI using the 638 nm laser line at 50% intensity. FRAP traces were normalized using the easyFRAP-web platform following the previously reported procedure^90^. Raw fluorescence intensities from the bleached ROI, the entire condensate containing the bleached ROI, and a background region were uploaded as ROI1, ROI2, and ROI3, respectively. Full-scale normalization was applied using the web platform, and the normalized fluorescence values were exported for further analysis. The normalized recovery curves were fitted in Origin using a double-exponential recovery function, 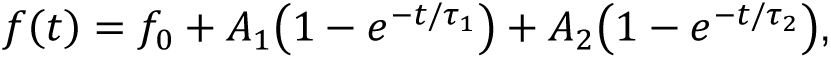 and the mobile fraction (MF) was calculated from the fitted recovery plateau as MF = *f*_O_ + *A*_1_ + *A*_2_ where *f*_O_ is the normalized intensity immediately after bleaching, *A*_1_ and *A*_2_ are the amplitudes of the fast and slow recovery components, and *τ*_1_ and *τ*_2_ are the corresponding relaxation times.

For whole-condensate FRAP measurements, the entire AF647-PRE condensate was selected as the bleach ROI, with an ROI area of approximately 2.5 µm². These measurements were acquired using the same AF647-PRE imaging and bleaching settings described above. Unlike the local FRAP measurements, whole-condensate recovery was analyzed from spatial intensity profiles. FRAP images were first processed with a Gaussian blur filter using a sigma of 0.5 µm, after which line intensity profiles were drawn across the condensate to quantify fluorescence recovery across the bleached droplet. Image acquisition and photobleaching were controlled using Leica LAS X software.

### Microfabrication and surface functionalization

The master wafer was prepared according to the previously described microfabrication method^91^, and the protocol was adjusted to attain a channel height of 20 µm^48^. The master wafer was placed on a flat support, and aluminium foil was wrapped around the wafer to form a shallow casting well. PDMS base and curing agent (SYLGARD^TM^ 184 elastomer) were mixed at a 10:1 mass ratio in a 50 mL tube using a spin rotor for 2 h. The mixed PDMS was poured over the wafer until the patterned features were fully covered and degassed in a desiccator until visible bubbles were removed. Excess mixed PDMS was stored at −20 °C and later used for coating glass slides. The PDMS was cured on the wafer for at least 3 h at 70 °C. After cooling, the cured PDMS slab was carefully peeled from the wafer, cut into individual device blocks using a sharp blade, and inlet and outlet openings were generated using appropriate biopsy punches.

For device bonding, glass coverslips were coated with a thin PDMS layer. A small volume of freshly mixed PDMS was dispensed onto the center of a clean glass coverslip and spread across the surface by gentle tilting. The coverslip was then mounted on a spin coater and coated using a two-step program: 30 s at 500 rpm with an acceleration of 300 rpm s^-1^, followed by 90 s at 1000 rpm with an acceleration of 1200 rpm s^-1^. The PDMS-coated coverslip was placed with the coated side facing upward on a raised support inside a covered dish and baked for 3 h at 70 °C to cure the coating. Before bonding, the channel side of each PDMS device block was cleaned with adhesive tape to remove dust and debris. The PDMS-coated glass surface was cleaned with pressurized air and kept with the coated side facing upward. Both surfaces were activated using an air plasma cleaner (Harrick Plasma PDC-32G) for 15 s at a radio frequency of 12 MHz. Immediately after plasma treatment, the PDMS device block was gently placed onto the PDMS-coated glass with the channel side facing down, taking care to avoid trapping air between the surfaces. The bonded devices were baked for approximately 2 h at 70 °C to strengthen the bond. The microfluidic devices were surface-functionalized with poly(vinyl alcohol) (PVA) to define the wetting properties required for double-emulsion production. The pressure controller was connected to an external compressed-air source and calibrated according to the manufacturer’s instructions. Four equal lengths of Tygon tubing were prepared for the three device inlets and for aspiration from the exit when required. The tubing was connected to reservoir holders using suitable microfluidic fittings, with each tube positioned at the bottom of its reservoir to prevent air from entering the lines during operation.

A 5% (w/v) PVA solution was prepared by dissolving PVA overnight at 80 °C with stirring and filtering the solution before use. After plasma bonding, devices were allowed to rest to permit partial relaxation of the plasma-induced hydrophilicity before selective PVA treatment. To functionalize the downstream channel region from the outer aqueous inlet to the production junction, a PVA-containing reservoir was connected to the outer aqueous inlet. Empty reservoirs were connected to the fluorinated oil and inner aqueous inlets. PVA was driven through the outer aqueous channel toward the outlet, while positive pressure was maintained on the inner aqueous and fluorinated oil channels to prevent PVA backflow into these regions. The treatment was continued for 5 min. To remove residual PVA, the PVA reservoir was disconnected from the outer aqueous inlet, the inner aqueous and fluorinated oil channels were flushed at 2 bar, and a vacuum of −950 mbar was simultaneously applied at the device exit to remove PVA from the exit region and outer aqueous inlet. Finally, the devices were baked for 15 min at 120 °C, allowed to cool, and stored at room temperature until use.

PDMS wells were prepared using clean, unpatterned silicon wafers as molds. PDMS base and curing agent were mixed as described above, poured over the silicon wafer, degassed, and cured at 70 °C. After curing, the PDMS layer was removed from the wafer and cut into individual blocks. Wells were generated by punching through the PDMS blocks using a 5 mm biopsy punch. The punched PDMS blocks were then plasma-bonded to PDMS-coated glass coverslips using the same bonding procedure described for the microfluidic devices. For experiments involving double-emulsion samples, the PDMS wells were treated with PVA. Briefly, the wells were filled with PVA solution and incubated for 15 min. The PVA solution was then removed by pipetting, and the bonded wells were baked on a hot plate at 120 °C for 15 min. For imaging condensates directly on glass, PDMS wells were instead treated with 1 mg mL^-1^ poly-L-lysine-grafted-polyethylene glycol (PLL-PEG) using the same procedure. After surface functionalization, all PDMS well devices were stored at 4 °C until use.

### Double emulsion production

Double emulsions (DEs) were generated using surface-functionalized microfluidic devices. Three phases were prepared before device operation: the inner aqueous phase (IA), the fluorinated oil phase (FO), and the outer aqueous phase (OA). The OA consisted of 1x PBS supplemented with 1% (v/v) Tween-20. The FO consisted of HFE-7500 fluorinated oil containing 2% (w/w) FluoSurf-C. The IA consisted of the required protein sample prepared in 1x PBS. The IA, FO, and OA solutions were loaded into separate reservoirs and connected to their corresponding device inlets using Tygon tubing. To initiate device operation, the OA was introduced first, followed by the FO, which allowed the formation of single emulsions at the first junction. The IA was then introduced to generate double emulsions at the downstream production junction. After confirming that no air bubbles were present in the channels, the inlet pressures were adjusted empirically until a stable and continuous DE pinch-off was obtained. Produced DEs were collected by inserting a clean 200 µL pipette tip into the device outlet. The collected DE dispersion was transferred into amber-colored bottles and stored at 4 °C until further use.

### pH-trigger experiments

Post-production pH-triggering experiments were performed in PDMS wells that had been surface-treated with 5% (w/v) PVA under the same conditions described above. The pH inside the double emulsions (DEs) was adjusted by exchanging the outer aqueous solution surrounding the DEs. For each experiment, 55 µL of an acidic feeding aqueous solution was pipetted into a PVA-treated PDMS well, followed by 5 µL of DE suspension. Either of the two acidic feeding solutions was used: pH 1 (100 mM HCl, 50 mM NaCl, 1% (v/v) Tween-20) and pH 2 (10 mM HCl, 140 mM NaCl, 1% (v/v) Tween-20). The DEs were then allowed to settle in the well. The well was kept covered with a coverslip throughout the experiment to minimize evaporation.

### Preparation of AcP/DiFMUP reaction samples

AcP-mediated hydrolysis of DiFMUP was used to monitor DiFMU formation in bulk solution and in the presence of PRE-based condensates. For microscopy experiments, samples containing 25 µM PRE and 100 µM DiFMUP were prepared in microcentrifuge tubes and incubated for 1 h before initiating the reaction by adding AcP to a final concentration of 0.05 U mL^-1^. AcP and DiFMUP were obtained from the EnzChek Acid Phosphatase Assay Kit (Thermo Fisher Scientific).

For microplate-reader assays, DiFMUP was used at final concentrations of 6.25, 12.5, 25, 37.5, 50, 62.5, 75, 87.5, and 100 µM. For the bulk condition, DiFMUP was incubated in buffer without PRE-based proteins. For the fusing-condensate condition, DiFMUP was incubated with 25 µM PRE. For the size-arrested-condensate condition, DiFMUP was incubated with 25 µM PRE and 1 µM Surf-PRE. All pre-reaction mixtures were prepared in microcentrifuge tubes and incubated for 1 h before being transferred to a Corning 384-well low-volume flat-bottom microplate. The final volume in each well did not exceed 40 µL. AcP was added last to initiate the reaction and was added in the following order for each experiment: bulk solution, fusing condensates, and size-arrested condensates.

### CLSM Imaging of AcP

Imaging of AF488-labeled AcP was performed on a Leica SP8 multiphoton microscope operated in confocal mode; multiphoton excitation was not used. Images were acquired in xyz mode using an HC PL IRAPO 40×/1.10 water-immersion objective with a zoom factor of 4. The pinhole was set to 78.7 µm, corresponding to 1.02 Airy units. Scanning was performed unidirectionally at 400 Hz without line or frame averaging. Image stacks were acquired at 1024 × 1024 pixels over a field of view of 72.66 × 72.66 µm, corresponding to a lateral pixel size of 0.071 µm. Z-stacks comprised 109 optical sections acquired at a step size of 0.43 µm, spanning a total axial distance of 46.44 µm. AcP fluorescence was excited using the 488 nm laser line at 10% intensity and detected between 499 and 560 nm using an internal photomultiplier tube (PMT) detector. Images were acquired using LAS X software.

### Multiphoton imaging of DiFMU fluorescence in condensates

Multiphoton fluorescence imaging was performed on a Leica SP8 multiphoton microscope to detect DiFMU fluorescence generated inside PRE condensates after the acid phosphatase (AcP)-catalyzed conversion of DiFMUP. AcP and DiFMUP were obtained from the EnzChek Acid Phosphatase Assay Kit (Thermo Fisher Scientific). A substrate-only control containing DiFMUP without AcP was imaged under the same acquisition conditions to assess background fluorescence and confirm that the signal increase required enzymatic product formation. Images were acquired using LAS X software in xyz mode with an HC PL IRAPO 40x/1.10 water-immersion objective. A single optical plane was recorded for each field-of-view using a 2104 x 2104 pixel format over a 290.62 x 290.62 µm field of view, corresponding to a pixel size of 0.138 µm. The scan was performed unidirectionally at 400 Hz with a zoom factor of 1. Frame averaging was set to 30, with no line averaging. DiFMU was excited using the multiphoton laser at 750 nm, with the laser intensity set to 1.0%. The visible laser lines at 488, 552, and 638 nm were switched off during acquisition. Fluorescence emission was collected using the HyD-RLD1 detector in counting mode over 412–511 nm, which covers the emission maximum of DiFMU (455 nm).

For time-lapse measurements of DiFMU accumulation, the AcP/DiFMUP reaction was imaged using the same multiphoton excitation and emission detection settings. Images were acquired in xyt mode using a 512 x 512 pixel format over a 72.66 x 72.66 µm field of view, corresponding to a pixel size of 0.142 µm. Time series were recorded for 15 frames over approximately 14 min, with a frame interval of approximately 60 s. These measurements were used to compare the temporal increase in DiFMU fluorescence within PRE condensates with that in the surrounding dilute phase.

### Microplate reader assay for DiFMU formation

DiFMU fluorescence was monitored using a BMG Labtech FLUOstar Omega microplate reader operated in fluorescence intensity plate mode. Fluorescence was measured from the top of the plate using an excitation filter of 340/10 nm and an emission filter of 450/10 nm. The settling time before each read was set to 0.5 s, and 20 flashes were used per measurement. Kinetic measurements were acquired for 300 cycles with a cycle time of 4 s. Between readings, the plate was shaken in double-orbital mode at 500 rpm to maintain sample homogeneity during the assay. Fluorescence intensity traces were recorded over time and used to compare DiFMU formation across the different reaction environments. AcP and DiFMUP were obtained from the EnzChek Acid Phosphatase Assay Kit (Thermo Fisher Scientific).

## ACKNOWLEDGEMENTS

We would like to thank Remco Fokkink for helping with pH measurements and GDL calibration curve, Rob de Haas for timely discussions on protein purification strategies, Ketan A. Ganar for discussions on strategies to improve microfabrication and the microfluidic production of double emulsions, Arjen Bader and Jan Willem Borst for training and assistance with advanced confocal microscopy, and Tijn Gaertner for assistance with the analysis of partitioning of fluorescently labeled AcP within condensates. We also thank Souma Majumdar, Thomas E. Kodger, and Manali Nandy for fruitful scientific discussions. S.D. acknowledges funding from the European Union’s Horizon Europe research and innovation programme under the Marie Skłodowska-Curie grant agreement No. 101119961 (SIGSYNCELL). The schematics were created with BioRender.com.

## AUTHOR CONTRIBUTIONS

U.G. and S.D. conceived the study and designed the experiments. U.G., E.v.d.V., and D.t.B. performed protein expression and purification, with methodological guidance from Z.H. U.G., Z.H., and C.Z. performed the MALDI-TOF experiments. U.G. and C.C. contributed to the methodology for microfabrication and double-emulsion production. U.G. performed all other experiments. U.G. and S.D. analyzed the data. U.G. and S.D. wrote the original draft. U.G., C.C., C.Z., R.d.V., J.v.d.G., and S.D. reviewed and edited the manuscript. S.D. was responsible for project administration and supervision. R.d.V. and J.v.d.G. provided additional scientific input through discussions of the results and project progress. All authors have read and approved the final version of the manuscript.

## DATA AVAILABILITY STATEMENT

Data supporting the findings of this study are available within the paper and its Supplementary Information. Any additional supporting data is available from the corresponding author upon reasonable request.

