## Supplementary Information for "Surface-stabilized sub-micron condensates for compartmentalizing synthetic cells and enhanced enzyme kinetics"

**for**

**Table of Contents:**

**Section 1:** Supplementary Tables (Page no. 3–4)

**Section 2:** Supplementary Note S1 (Page no. 5–7)

**Section 3:** Figure S1-S10 (Page no. 8–19)

**Section 4:** Caption for Supplementary Video S1 (Page no. 20)

### Supplementary Tables

**Table S1.** PRE and Surf-PRE sequences. The PRE sequences were designed with a C-terminal WPC tag, in which tryptophan (W) enables protein quantification by absorbance at 280 nm, proline (P) serves as a spacer, and cysteine (C) provides a thiol group for maleimide-based fluorescent labelling.

| **Protein** | **Sequence** |
|---|---|
| PRE | MGVGVPGYGVPGEGVPGVGVPGYGVPGVGVPGEGVPGYGVPGVG<br>VPGYGVPGHGVPGYGVPGVGVPGEGVPGYGVPGVGVPGYGVPGE<br>GVPGVGVPGYGVPGVGVPGYGVPGEGVPGVGVPGYGVPGVGVPG<br>EGVPGYGVPGVGVPGYGVPGHGVPGYGVPGVGVPGEGVPGYGVP<br>GVGVPGYGVPGEGVPGVGVPGYGVPGVGVPGYGVPGEGVPGVGV<br>PGYGVPGVGVPGEGVPGYGVPGVGVPGYGVPGHGVPGYGVPGVG<br>VPGEGVPGYGVPGVGVPGYGVPGEGVPGVGVPGYGVPGWPC |
| Surf-PRE | MGSGVPGSGVPGSGVPGSGVPGSGVPGSGVPGSGVPGSGVPGSG<br>VPGSGVPGSGVPGSGVPGSGVPGSGVPGSGVPGSGVPGSGVPGS<br>GVPGSGVPGSGVPGSGVPGSGVPGSGVPGSGVPGSGVPGSGVPG<br>SGVPGSGVPGSGVPGSGVPGSGVPGSGVPGSGVPGSGVPGSGVP<br>GSGVPGSGVPGSGVPGSGVPGSGVPGSGVPGSGVPGSGVPGSGV<br>PGSGVPGSGVPGSGVPGSGVPGSGVPGSGVPGSGVPGSGVPGSG<br>VPGSGVPGSGVPGSGVPGSGVPGSGVPGSGVPGSGVPGVGVPGY<br>GVPGEGVPGVGVPGYGVPGVGVPGEGVPGYGVPGVGVPGYGVPG<br>HGVPGYGVPGVGVPGEGVPGYGVPGVGVPGYGVPGEGVPGVGVP<br>GYGVPGVGVPGYGVPGEGVPGVGVPGYGVPGVGVPGEGVPGYGV<br>PGVGVPGYGVPGHGVPGYGVPGVGVPGEGVPGYGVPGVGVPGYG<br>VPGEGVPGVGVPGYGVPGVGVPGYGVPGEGVPGVGVPGYGVPGV<br>GVPGEGVPGYGVPGVGVPGYGVPGHGVPGYGVPGVGVPGEGVPG<br>YGVPGVGVPGYGVPGEGVPGVGVPGYGVPGWPC |

**Table S2.** List of sequencing primers used in this study.

| **Primers** | **5' to 3' sequences** |
|---|---|
| Forward | TAATACGACTCACTATAGGG |
| Reverse | GCTAGTTATTGCTCAGCGG |

**Table S3.** Michaelis-Menten kinetic parameters for acid phosphatase-catalyzed hydrolysis of DiFMUP in different reaction environments. Apparent $V_{max}$, $K_m$, and catalytic efficiency

($V_{max}/K_m$) values were obtained from Michaelis–Menten fits to the initial reaction phase and the later eye-like fluorescence phase for reactions performed in bulk solution (DiFMUP + AcP), PRE MOs (PRE + DiFMUP + AcP), and Surf-PRE-stabilized MOs (PRE + Surf-PRE + DiFMUP + AcP).

| **Reaction Condition** | **$V_{max}$ (μM/s)** | | **$K_m$ (μM)** | | **Apparent catalytic efficiency ($s^{-1}$) ($V_{max}/K_m$)** | |
|---|---|---|---|---|---|---|
| | Regime I | Regime II | Regime I | Regime II | Regime I | Regime II |
| DiFMUP + AcP | 26 | 30 | 84 | 81 | 0.31 | 0.37 |
| PRE + DiFMUP + AcP | 55 | 100 | 145 | 220 | 0.38 | 0.45 |
| PRE + Surf-PRE + DiFMUP + AcP | 67 | 352 | 133 | 570 | 0.5 | 0.62 |

**Note S1.** Parameters used for estimating the condensate volume fraction and PRE distribution within double emulsions. Experimental and molecular parameters used to calculate the fraction of the double-emulsion volume occupied by PRE condensates and to estimate the distribution of PRE between the condensed and dilute phases.

| **Parameter(s)** | **Value used** |
|---|---|
| PRE concentration, $C_{\mathrm{PRE,\,T}}$ | 20 µM |
| Surf-PRE concentration, $C_{\mathrm{Surf\text{-}PRE,\,T}}$ | 0.1 µM |
| Double emulsion diameter | 31.5 µm |
| Double emulsion radius, $r_{\mathrm{DE}}$ | 15.8 µm |
| Average number of condensates per DE, $n_{\mathrm{C}}$ | 5.3 (~ 5) |
| Average condensate radius in DE, $r_{\mathrm{C}}$ | 0.9 µm |
| PRE radius of gyration, $r_{\mathrm{g,\,PRE}}$ | 6.0 nm |
| PRE dense-phase partition coefficient, $K_{\mathrm{PRE,\,C}}$ | 454.5 |

Geometric estimation of the condensate volume fraction and partitioning-based calculation of PRE distribution within double emulsions. The measured double-emulsion and condensate dimensions were used to estimate the total volume occupied by PRE condensates and the corresponding condensate volume fraction within each double emulsion. The experimentally determined PRE partition coefficient was then combined with this volume fraction and a mass-balance expression to estimate the PRE concentrations in the condensed and dilute phases, as well as the fraction of the total PRE present in each phase.

$$\text{Volume occupied by each DE, } V_{\mathrm{DE}} = \frac{4}{3}\pi \times (15.8)^3 = 16522\ \mu m^3 \approx 1.7 \times 10^{-11}\ L$$

$$\text{Total volume occupied by condensates, } V_{\mathrm{PRE,\,C}} = n_{\mathrm{C}} \times \text{Volume occupied by one condensate}$$

$$= 5 \times \frac{4}{3}\pi \times (0.9)^3 = 15.3\ \mu m^3$$

$$\text{Volume fraction of condensates, } \phi_{\mathrm{C}} = \frac{V_{\mathrm{PRE,\,C}}}{V_{\mathrm{DE}}} = \frac{15.3 \times 10^{-15}}{1.7\ \times 10^{-11}} = 9 \times 10^{-4} = 0.0009\ \approx 0.1\%$$

Therefore, PRE condensates occupy only 0.1% of the DE volume.

$$\text{Partition coefficient of PRE in dense phase as compared to dilute phase, } K_{\mathrm{PRE}} = \frac{C_{\mathrm{PRE,\,C}}}{C_{\mathrm{PRE,\,D}}}$$

$$\Rightarrow C_{\mathrm{PRE,\,C}} = K_{\mathrm{PRE}} C_{\mathrm{PRE,\,D}}$$

From the mass-balance expression,

$$C_{\text{PRE, T}} = C_{\text{PRE, C}}\phi_{\text{C}} + C_{\text{PRE, D}}(1 - \phi_{\text{C}})$$

$$\Rightarrow C_{\text{PRE, T}} = K_{\text{PRE}}C_{\text{PRE, D}}\phi_{\text{C}} + C_{\text{PRE, D}}(1 - \phi_{\text{C}})$$

$$\Rightarrow C_{\text{PRE, T}} = C_{\text{PRE, D}}(K_{\text{PRE}}\phi_{\text{C}} + (1 - \phi_{\text{C}}))$$

$$\Rightarrow C_{\text{PRE, D}} = \frac{C_{\text{PRE, T}}}{K_{\text{PRE}}\phi_{\text{C}} + (1 - \phi_{\text{C}})}$$

Therefore,

$$C_{\text{PRE, D}} = \frac{20}{454.5 \times 0.0009 + (1 - 0.0009)} = 14.2\ \mu M$$

$$C_{\text{PRE, C}} = K_{\text{PRE}}C_{\text{PRE, D}} = 6455.3\ \mu M = 6.5\ mM$$

Now,

$$\phi_{\text{C}} = \frac{V_{\text{PRE, C}}}{V_{\text{DE}}} \Rightarrow V_{\text{PRE, C}} = \phi_{\text{C}}V_{\text{DE}}$$

$$V_{\text{PRE, D}} = (1 - \phi_{\text{C}})V_{\text{DE}}$$

Here,

$$n_{\text{PRE, T}} = n_{\text{PRE, C}} + n_{\text{PRE, D}}$$

$$\Rightarrow n_{\text{PRE, T}} = C_{\text{PRE, C}}V_{\text{PRE, C}} + C_{\text{PRE, D}}V_{\text{PRE, D}}$$

$$\Rightarrow n_{\text{PRE, T}} = C_{\text{PRE, C}}\phi_{\text{C}}V_{\text{DE}} + C_{\text{PRE, D}}(1 - \phi_{\text{C}})V_{\text{DE}}$$

$$\Rightarrow n_{\text{PRE, T}} = K_{\text{PRE}}C_{\text{PRE, D}}\phi_{\text{C}}V_{\text{DE}} + C_{\text{PRE, D}}(1 - \phi_{\text{C}})V_{\text{DE}}$$

$$\Rightarrow n_{\text{PRE, T}} = C_{\text{PRE, D}}V_{\text{DE}}(K_{\text{PRE}}\phi_{\text{C}} + (1 - \phi_{\text{C}}))$$

Now,

$$\text{Fraction of total PRE in the dense phase, } \chi_{\text{PRE, C}} = \frac{n_{\text{PRE, C}}}{n_{\text{PRE, T}}}$$

$$\Rightarrow \chi_{\text{PRE, C}} = \frac{K_{\text{PRE}}C_{\text{PRE, D}}\phi_{\text{C}}V_{\text{DE}}}{C_{\text{PRE, D}}V_{\text{DE}}(K_{\text{PRE}}\phi_{\text{C}} + (1 - \phi_{\text{C}}))}$$

$$\Rightarrow \chi_{\text{PRE, C}} = \frac{K_{\text{PRE}}\phi_{\text{C}}}{K_{\text{PRE}}\phi_{\text{C}} + (1 - \phi_{\text{C}})}$$

$$\Rightarrow \chi_{\text{PRE, C}} = \frac{454.5 \times 0.0009}{454.5 \times 0.0009 + (1 - 0.0009)} = 0.290 \approx 29.0\%$$

Therefore, 29.0% of PRE is in the dense condensate phase, and 71.0% of PRE is in the dilute phase.

**Supplementary Figures**

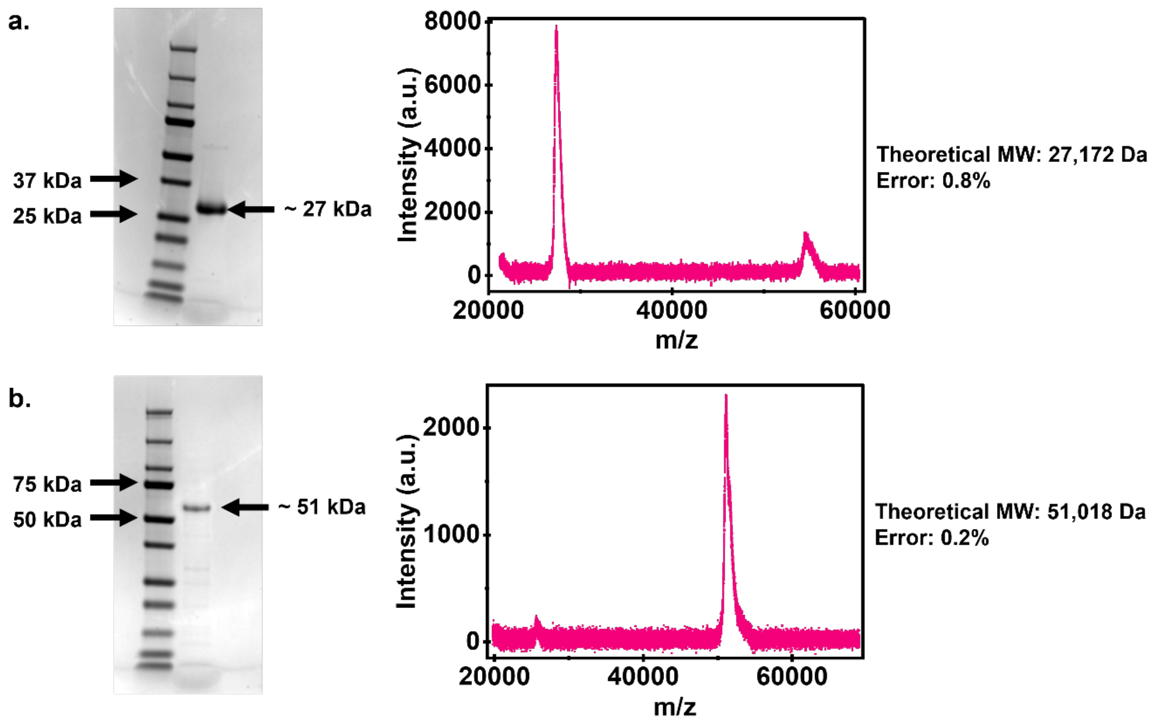


**Figure S1. Expression, purification, and molecular-weight characterization of PRE and Surf-PRE.** a) SDS-PAGE analysis of PRE following recombinant expression and purification by inverse transition cycling (ITC), with the purified protein shown in the middle lane. The corresponding MALDI-TOF mass spectrum shows a dominant peak at the characteristic mass-to-charge ratio (m/z) of PRE, assigned to the singly charged molecular ion, for which the m/z value closely approximates the molecular mass in Da. b) SDS–PAGE analysis of Surf-PRE following recombinant expression and purification by ITC, together with the corresponding MALDI-TOF mass spectrum showing its dominant m/z peak. The deviations of the experimentally determined molecular weights, derived from the measured m/z values, from the theoretical molecular weights of PRE and Surf-PRE are indicated.

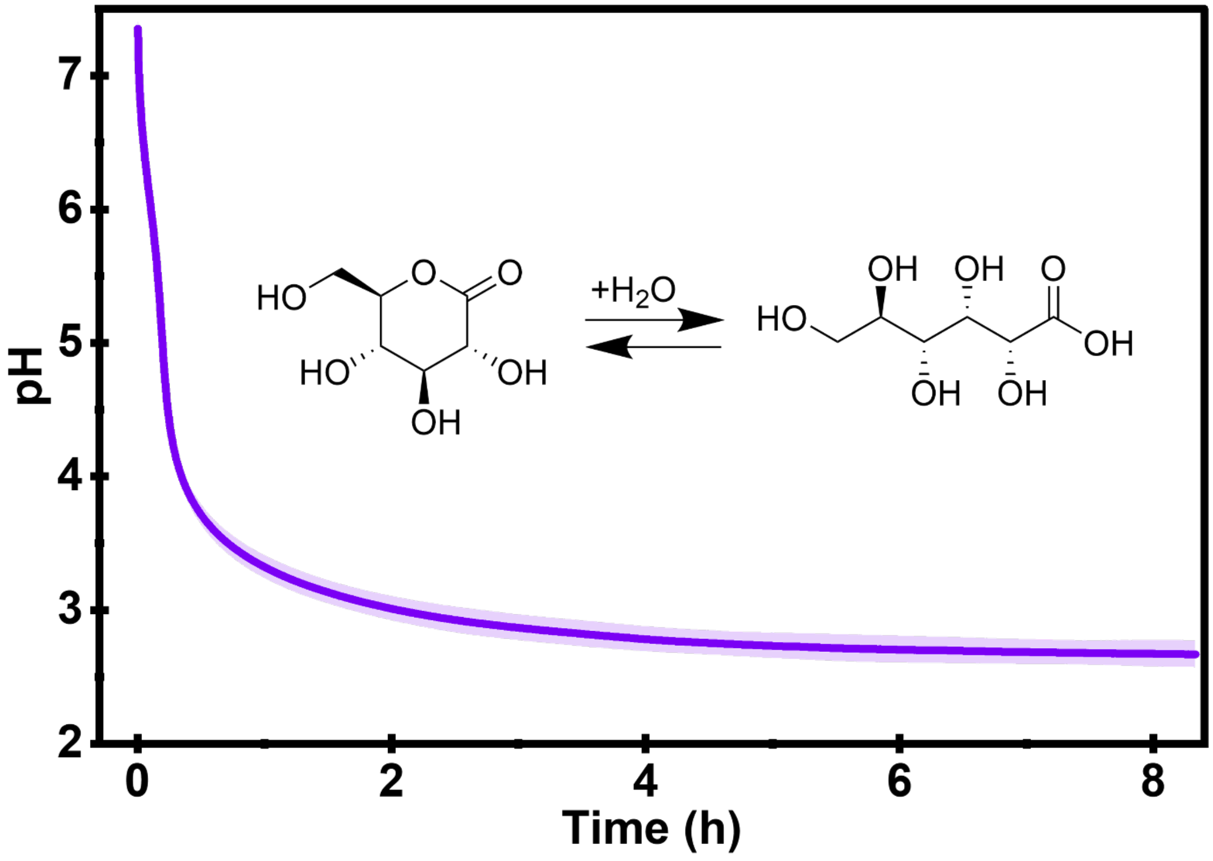


**Figure S2. Calibration of GDL-mediated acidification in PBS.** Time-dependent pH evolution following the addition of 84.2 mM glucono-δ-lactone (GDL) to PBS. Hydrolysis of GDL induces a rapid initial decrease in pH, followed by a slower acidification phase and gradual approach toward a plateau. Data are presented as mean ± s.d., represented by shaded areas (n = 3).

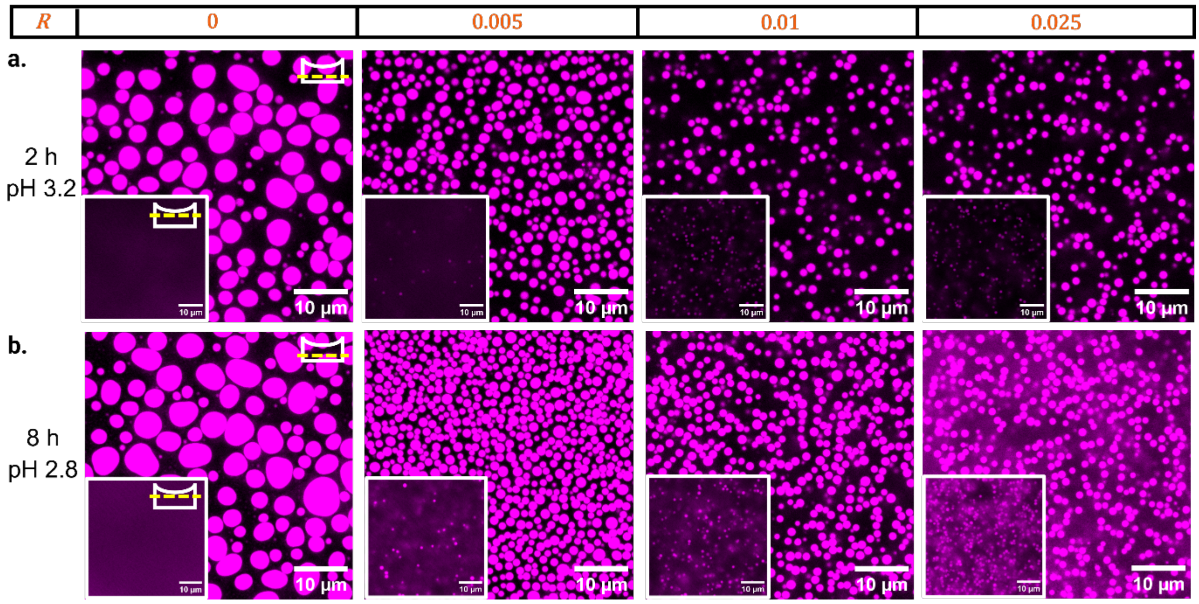


**Figure S3. Gradual acidification by GDL results in a less pronounced Surf-PRE-dependent regulation of PRE condensate size.** a,b) Fluorescence microscopy images of PRE condensates formed at different *R* values, 2 h (a) and 8 h (b) after the GDL trigger. Insets show the field-of-view in the bulk of the solution to indicate the presence of dispersed condensates. Increasing *R* results in smaller condensates at both time points; however, the change in condensate size is less pronounced than that observed following the sharp 50 mM HCl trigger (pH 1.3).

.

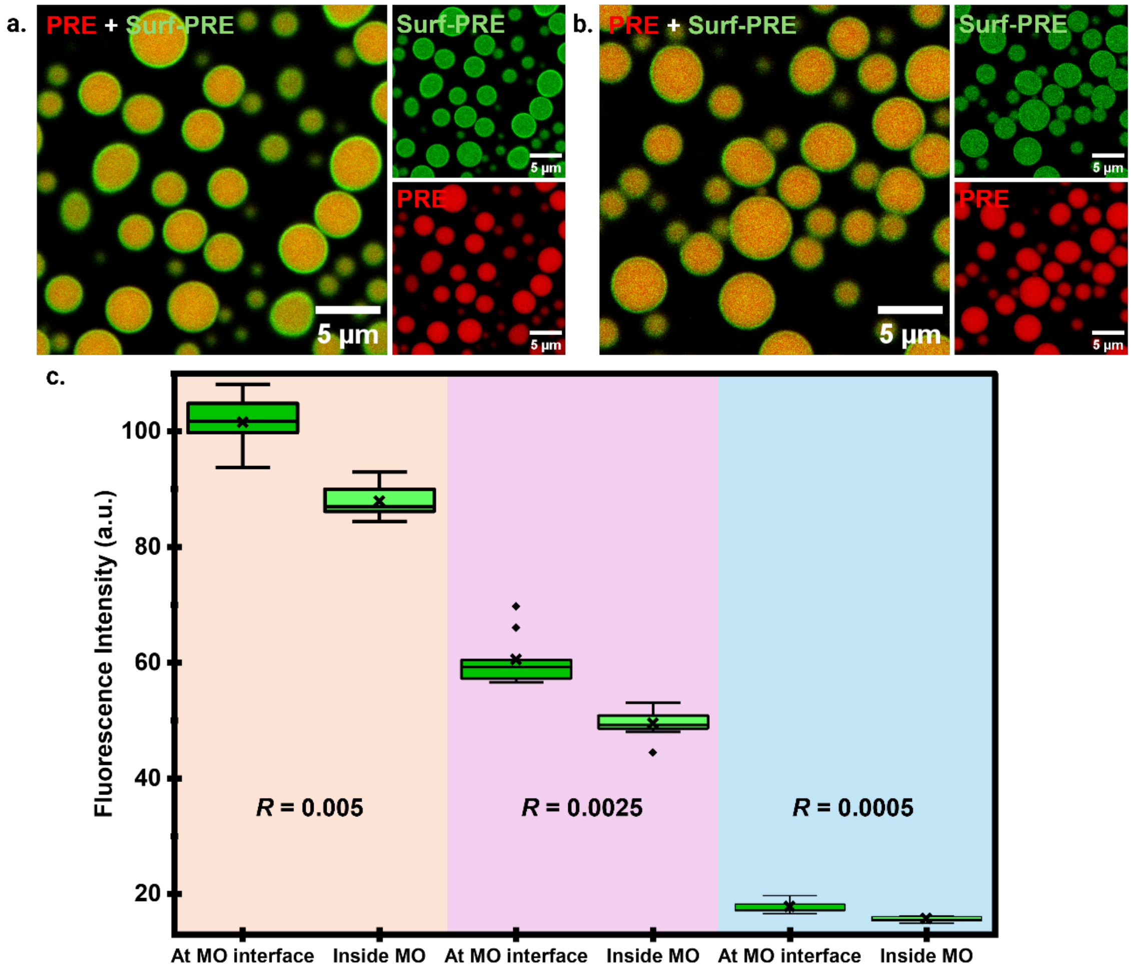


**Figure S4. Relative distribution of Surf-PRE between the MO interface and interior at different Surf-PRE concentrations.** (a,b) Representative CLSM images of MOs formed at *R* = 0.0025 and *R* = 0.0005, respectively. AF647-PRE is shown in red and AF488-Surf-PRE in green. (c) Box-and-whisker plots of the raw AF488-Surf-PRE fluorescence intensities measured at the MO interface and within the MO interior for *R* = 0.005, 0.0025, and 0.0005. Boxes represent the interquartile range, central lines indicate the median, crosses indicate the mean, and diamonds indicate outliers. The corresponding interface-to-interior intensity ratios were approximately 1.2, 1.2, and 1.1, respectively.

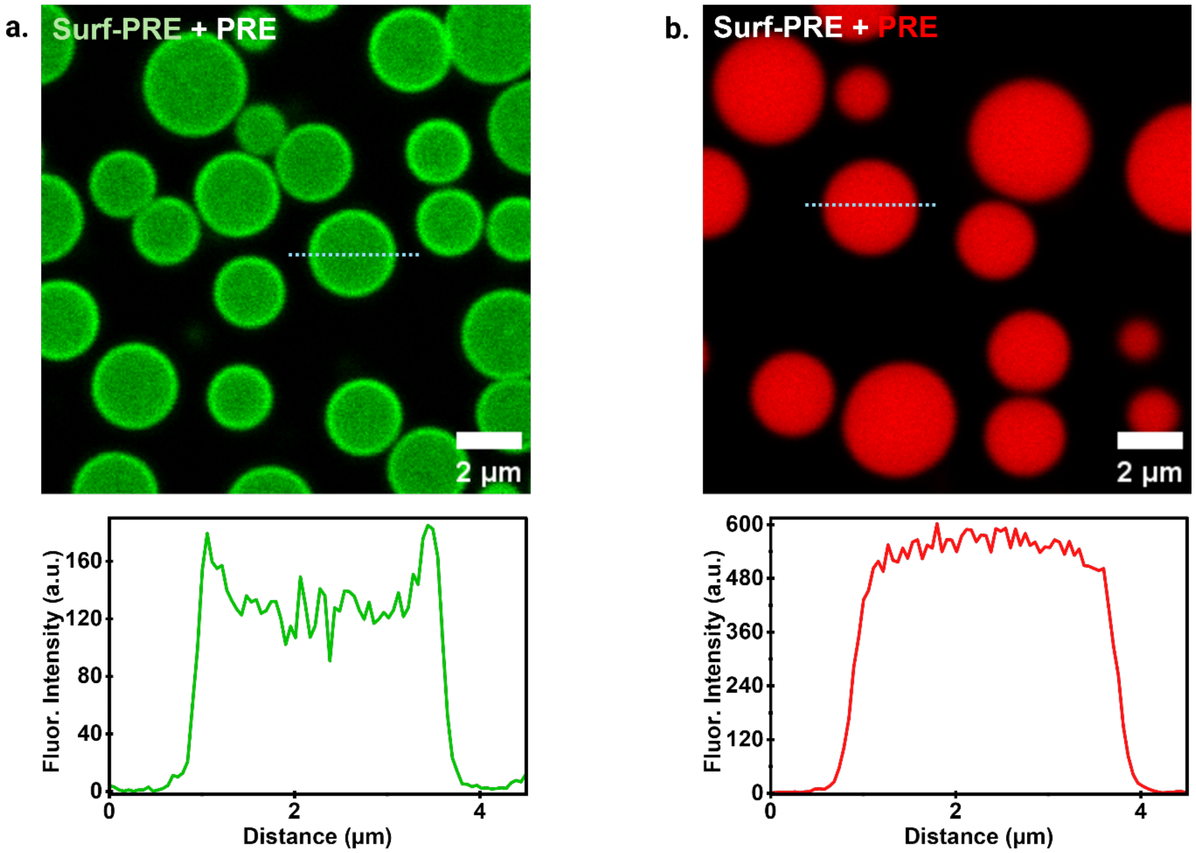


**Figure S5. Single-channel controls confirming the spatial distributions of PRE and Surf-PRE within MOs.** (a) MOs containing AF488-Surf-PRE and unlabeled PRE show Surf-PRE within the condensate interior together with enrichment at the interface, whereas (b) MOs containing AF647-PRE and unlabeled Surf-PRE show homogeneous PRE distribution without interfacial enrichment.

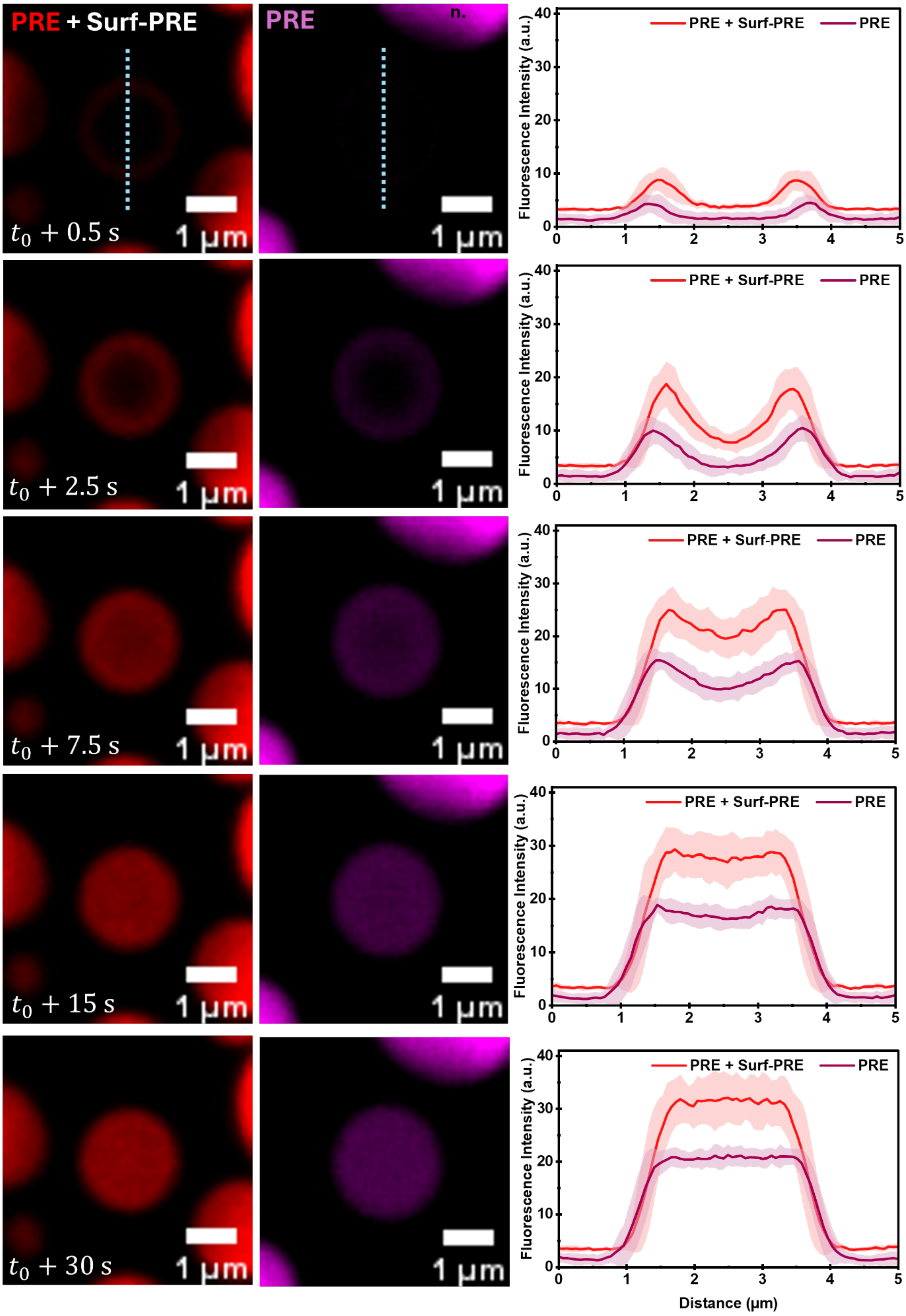
PRE + Surf-PRE
PRE
$t_0 + 0.5$ s
$t_0 + 2.5$ s
$t_0 + 7.5$ s
$t_0 + 15$ s
$t_0 + 30$ s
1 µm
Fluorescence Intensity (a.u.)
Distance (µm)

**Figure S6. Whole-droplet FRAP reveals inward recovery of PRE from the MO boundary.** (a,b) Representative time-lapse CLSM images showing AF647-PRE recovery after bleaching the entire MO in Surf-PRE-stabilized MOs and PRE-only MOs, respectively. Images were acquired at 0.5, 2.5, 7.5, 15, and 30 s after bleaching. Cyan dotted lines indicate the positions used to obtain the fluorescence intensity profiles. (c) Corresponding line intensity profiles across the bleached MOs at each time point. Fluorescence initially reappeared at the MO periphery and subsequently progressed toward the interior, consistent with molecular influx from the surrounding dilute phase. PRE-only MOs showed a slower inward recovery than Surf-PRE-stabilized MOs.

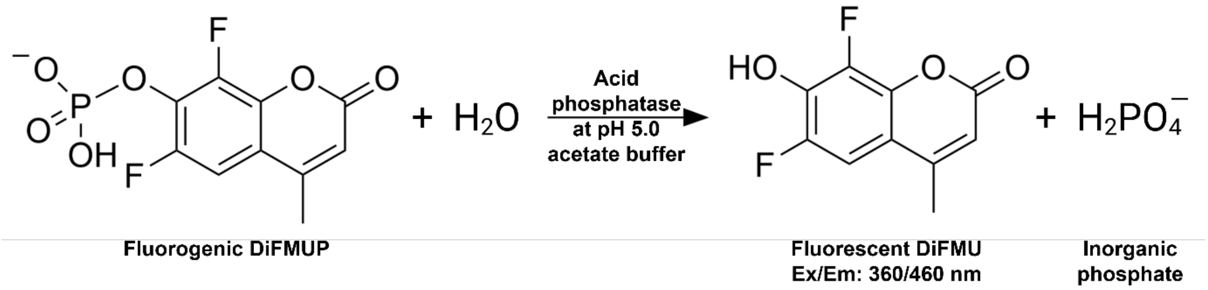


**Figure S7. Acid phosphatase-catalyzed hydrolysis of DiFMUP.** Schematic representation of the dephosphorylation of 6,8-difluoro-4-methylumbelliferyl phosphate (DiFMUP) to the fluorescent product 6,8-difluoro-4-methylumbelliferone (DiFMU) by acid phosphatase in acetate buffer at pH 5.0.

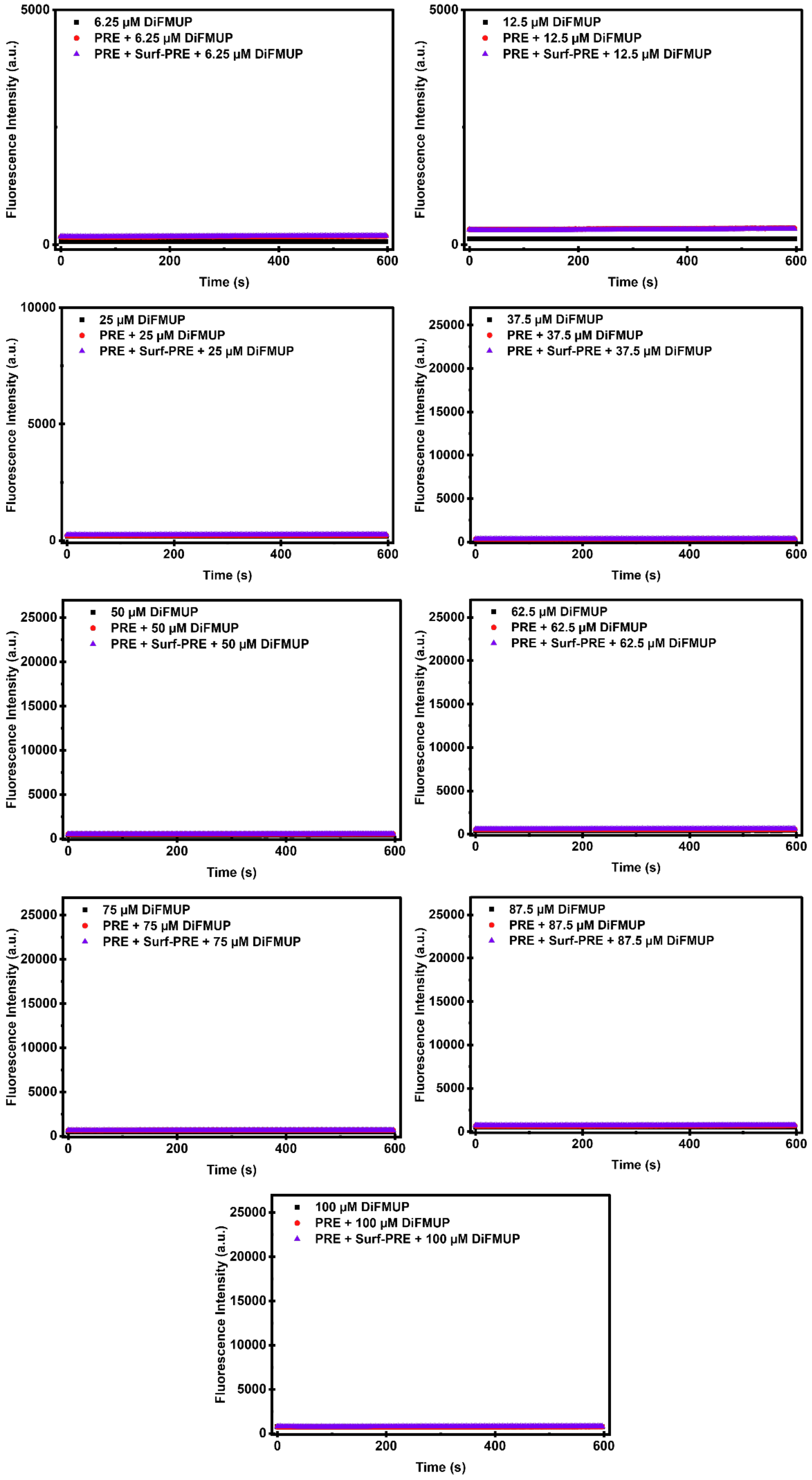

6.25 μM DiFMUP
PRE + 6.25 μM DiFMUP
PRE + Surf-PRE + 6.25 μM DiFMUP
12.5 μM DiFMUP
PRE + 12.5 μM DiFMUP
PRE + Surf-PRE + 12.5 μM DiFMUP
25 μM DiFMUP
PRE + 25 μM DiFMUP
PRE + Surf-PRE + 25 μM DiFMUP
37.5 μM DiFMUP
PRE + 37.5 μM DiFMUP
PRE + Surf-PRE + 37.5 μM DiFMUP
50 μM DiFMUP
PRE + 50 μM DiFMUP
PRE + Surf-PRE + 50 μM DiFMUP
62.5 μM DiFMUP
PRE + 62.5 μM DiFMUP
PRE + Surf-PRE + 62.5 μM DiFMUP
75 μM DiFMUP
PRE + 75 μM DiFMUP
PRE + Surf-PRE + 75 μM DiFMUP
87.5 μM DiFMUP
PRE + 87.5 μM DiFMUP
PRE + Surf-PRE + 87.5 μM DiFMUP
100 μM DiFMUP
PRE + 100 μM DiFMUP
PRE + Surf-PRE + 100 μM DiFMUP
Fluorescence Intensity (a.u.)
Time (s)
0
200
400
600
5000
10000
15000
20000
25000

**Figure S8. Control measurements in the absence of acid phosphatase.** Fluorescence intensity was monitored for 600 s at DiFMUP concentrations ranging from 6.25–100 μM in bulk solution, in the presence of 25 μM PRE, and in the presence of 25 μM PRE + 1 μM Surf-PRE. No increase in fluorescence was observed under any condition, confirming that DiFMU formation in the corresponding reaction experiments resulted from acid phosphatase-catalyzed DiFMUP hydrolysis. Data are presented as mean ± s.d., represented by shaded areas (n = 3 independent bulk reactions for each condition).

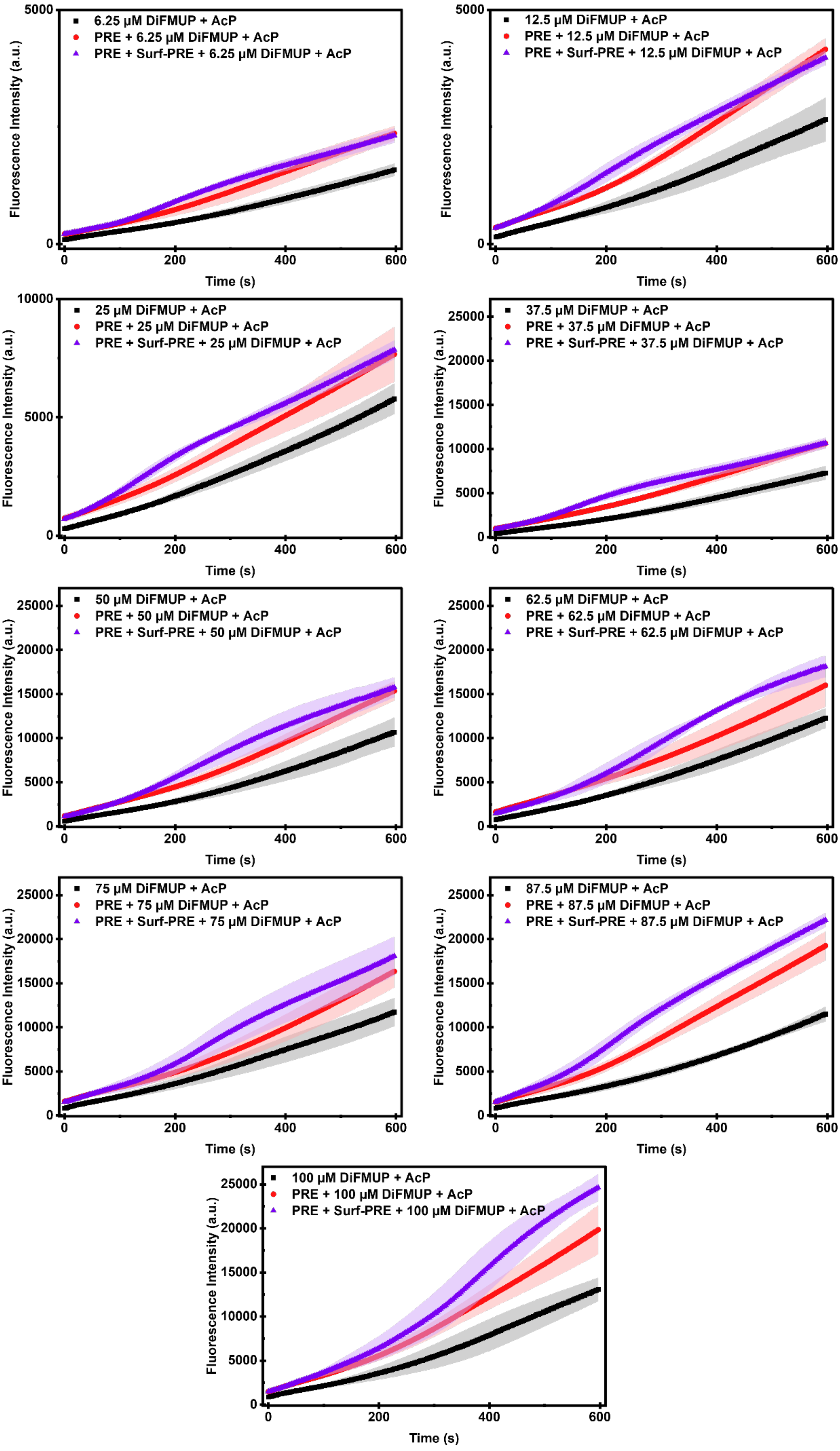

6.25 μM DiFMUP + AcP
PRE + 6.25 μM DiFMUP + AcP
PRE + Surf-PRE + 6.25 μM DiFMUP + AcP
12.5 μM DiFMUP + AcP
PRE + 12.5 μM DiFMUP + AcP
PRE + Surf-PRE + 12.5 μM DiFMUP + AcP
25 μM DiFMUP + AcP
PRE + 25 μM DiFMUP + AcP
PRE + Surf-PRE + 25 μM DiFMUP + AcP
37.5 μM DiFMUP + AcP
PRE + 37.5 μM DiFMUP + AcP
PRE + Surf-PRE + 37.5 μM DiFMUP + AcP
50 μM DiFMUP + AcP
PRE + 50 μM DiFMUP + AcP
PRE + Surf-PRE + 50 μM DiFMUP + AcP
62.5 μM DiFMUP + AcP
PRE + 62.5 μM DiFMUP + AcP
PRE + Surf-PRE + 62.5 μM DiFMUP + AcP
75 μM DiFMUP + AcP
PRE + 75 μM DiFMUP + AcP
PRE + Surf-PRE + 75 μM DiFMUP + AcP
87.5 μM DiFMUP + AcP
PRE + 87.5 μM DiFMUP + AcP
PRE + Surf-PRE + 87.5 μM DiFMUP + AcP
100 μM DiFMUP + AcP
PRE + 100 μM DiFMUP + AcP
PRE + Surf-PRE + 100 μM DiFMUP + AcP
Fluorescence Intensity (a.u.)
Time (s)

**Figure S9. Acid phosphatase-catalyzed hydrolysis of DiFMUP under different reaction conditions.** Time-dependent DiFMU fluorescence was monitored for 600 s at DiFMUP concentrations of 6.25–100 μM. Reactions were performed in bulk solution containing DiFMUP and acid phosphatase (black), in the presence of 25 μM PRE (red), or in the presence of 25 μM PRE and 1 μM Surf-PRE (purple). Data are presented as mean ± s.d., represented by shaded areas (n = 5 independent bulk reactions for each condition).

**Caption for Supplementary Video**

**Supplementary Video S1. Formation and persistence of multiple MOs in synthetic cells during acidification.** Representative fluorescence time-lapse showing the emergence and persistence of multiple MOs in DEs containing 20 μM PRE, including 4 mol% AF647-PRE, and 2 μM Surf-PRE during a fast acidification trigger (100 mM HCl, 50 mM NaCl, and 1% Tween-20).
